# A mathematical model for Dengue and Chikungunya in Mexico

**DOI:** 10.1101/122556

**Authors:** Nohemí Báez-Hernández, Mauricio Casas-Martínez, Rogelio Danis-Lozano, Jorge X. Velasco-Hernández

## Abstract

We present a model that incorporates two co-circulating viral diseases, Dengue and Chikungunya, where we allow secondary infections from either of the two diseases. We only consider one vector population, *Ae. aegypti* since in the Mexican region where we set our scenarios, only this species has been reported to transmit both viruses. We estimate the basic reproduction number and perform numerical simulations for different scenarios where we may observe coexistence of Dengue and Chikungunya; we also compare the results of the model with Dengue and Chikungunya data from Mexico 2015 and we obtain a good model fit. To complete our findings we perform a sensitivity analysis, and calculate the partial rank correlation coefficients (PRCCs) to determine the parameter values influence on the reproduction numbers and predict fate of the diseases.

We show that *R*_0_ for each one of the viruses is highly sensitive to the mosquito biting rate and the transmission rates for both diseases with positive influence and the average lifespan of mosquito along with the human recovery rate with negative influence on both diseases. Our results are consistent with those of previous authors.

## 1 Introduction

Chikungunya and Dengue fever are mosquito-borne infectious diseases that are a publie health problem of considerable importance, particularly throughout the American Continent.

Dengue occurs in tropical and subtropical areas and can infect millions of people every year, and now it has increased on climate change [1]. In recent years, Chikungunya virus re-emerged and caused outbreaks in Italy, Asia and several Indian Ocean islands [1].

Dengue modelling is more frequent and there are more strategies for its control, in contrast to Chikungunya virus. Chikungunya attracted worldwide attention, in 2015 particularly in Mexico. The main vectors for both diseases are *Ae. aegypti* and *Ae. albopictus* [1], the last one is a more efficient vector in the transmission of CHIKV-R (a new 2005 Reunion strain of Chikungunya) than it is for Dengue [1].

The principal vector of Dengue transmission in the Americas is *Ae. aegypti* [1]. Dengue causes a range of illnesses in humans, from asymptomatic cases to severe disease, sometimes fatal [2], and generally occurs in older children and adults. Among the most common symptoms are: an abrupt onset of fever, frontal headache, nausea, vomiting and others. [2]. The acute phase of the disease lasts 3 to 7 days [2]. The two most severe forms of dengue: hemorrhagic fever (DHF) and its associated Dengue Shock Syndrome (DSF) usually occur in children under 15 years, although these can also occur in adults [2]. There are four Dengue serotypes; the infection by any of their strains produces long term immunity, but only a short period of cross-immunity to other three serotypes [2]. The extrinsic incubation period lasts between 8 to 14 days after the mosquito eats infected blood. The mosquito remains infectious for life [2]. In endemic tropical areas, Dengue is spread throughout the year, but with a cyclical pattern. The severity of Dengue is associated with secondary infections, although its causes have not been explained [2]. There are many mitigation strategies for Dengue, for example: indoors and outdoors spraying (adulticides, larvicides), lethal ovitraps [1], removing artificial oviposition sites (tins, tires, flowerpot, etc.), all these for reduction of the mosquito population; also the use of screens or mosquito repellent for the decrease of human exposure to mosquito bites, vaccination, among others. However, these are not always effective and, as a result, Dengue prevalence has increased in the last 40 years [2], recent studies indicate about 390 million Dengue infected appear annually [3].

Chikungunya fever (CHIKF) is a mosquito-borne viral infection, which it was first isolated in Tanzania in 1953 [4]. Chikungunya is frequently confused with Dengue. One specific symptom to Chikungunya is a debilitating and prolonged joint pain, where the peripheral small joints are the most affected [4]; other symptoms that most patients present are: swelling and muscle stiffness [42]. Chikungunya virus has an intrinsic incubation period in the human host of about 4 days after an infected mosquito bite, and the illness lasts a period of 7 days approximately. It is during this period that mosquitoes can be infected with CHIKV when biting infected hosts [4]. Chikungunya infected individuals exhibit a severe joint pain that can last for a long time after the acute phase of the disease, until 3 years [42]. There is evidence showing that 25% of infected individuals suffer mild symptoms or are asymptomatic [4]. Once infected the mosquitoes remain infectious for life [4].

It is necessary to understand how both diseases are spread and persist and which are the most important parameters involved in their dynamics, that may help to develop strategies for prevention, control, and joint treatments. There are many models for Dengue in the literature e.g. [2, 5, 6, 7, 44], some others for Chikungunya [8, 9, 10, 25, 43, 46] that explore different aspects of their spread and behavior; more recently some works are emerged, which study both diseases e.g [45, 47]. Unlike previous models, we model the dynamics of both diseases simultaneously since we couple them.

Chikungunya received attention after several outbreaks from 2013 to the date, these have occurred in places where Dengue fever is endemic. Around of 154 and 99 countries in the world have reported endemic or epidemic Dengue and Chikungunya, respectively; moreover, 98 places notified transmission of both viruses, of which only 13 registered co-infections [3]. For example, the transmission in the Caribbean island of Saint Martin in 2013 took place together with a Dengue epidemic where sixteen coinfected cases were documented [3]; other example is Delhi, India an area with both viruses co-circulating and where *Ae. aegypti* mosquitoes are present, co-infections were detected during a outbreak of Dengue in 2006 [11].

The etiologic strain of Chikungunya transmission occurred in the Americas was an Asian lineage [8, 9]. After quickly sweeping through the Caribbean following its initial discovery in St. Martin, this Asian CHIKV strain has now spread into all Central American countries, most of South America, and into northern Mexico. Until 2016, in this continent, 44 countries have reported Chikungunya infected [3].

In 2014, the Soconusco region (Southern Chiapas), showed a major outbreak concentrating most of the cases in the cities where the record of autochthonous cases of Chikungunya occurred. This entomological scenario allowed the establishment of local highly competent mosquito populations to receive and spread CHIKV. The first autochthonous case of Chikungunya fever in Mexico was officially notified on November 7th, 2014 in Arriaga, Chiapas, 300 km from the border with Guatemala [12]. The first autochthonous cases were reported in other states: Campeche, Colima, Chiapas, Guerrero, Michoacán, Morelos, Oaxaca, Tabasco and Veracruz. The state of Jalisco reported one single case [13]. The epidemiological situation described above occurred during the dry season when the densities of *Aedes* mosquitoes were low. For 2015, more that 11500 cases was registered throughout the country, the states with the highest number of cases were: Guerrero, Veracruz and Yucatán [14].

In the Mexican region where we set up our scenarios only *Ae. aegypti* has been reported to transmit both viruses, and therefore we postulate only one type of vector population. We allow secondary infections from either of the two diseases. We use a *SEIR* model for the dynamics of Dengue and Chikungunya coupled with a *SI* model that describes the dynamic of the vector population. For Chikungunya there are reports that after 5 to 7 days (duration of its acute phase, *I_c_* in our model, see Section 2) around of 50% of the infected individuals are recovered, and the other half may develop a sub-acute phase [41], which we name it *I*_2*c*_, where joint pain can occur in the 2 or 3 months following with the same intensity as the acute phase; then the infected individuals pass to the chronic phase, *I*_3*c*_ for us and the last phase of this disease [40], here the patient can present joint pain in the 6, 12, 18 or 24 months following. In these two last phases the infected individuals have some degree of clinical deterioration that weaken them and make them more susceptible to other infections, particularly to Dengue infection. Note that the infected individual in *I*_2c_ or *I*_3c_ can not be recovered. We consider that this categorization of Chikugunya disease is important to explore the susceptibility to Dengue disease, which is of interest for us.

We assume that there is vertical transmission of the Dengue disease in vector population, i.e., if *q* is the recruitment rate of mosquitoes, a fraction *qp_d_* are born infectious and a fraction *q*(1 – *p_d_*) are born susceptible. Studies in Mexico show evidence of the existence of vertical transmission of Dengue in *Ae. aegypti*, whose importance lies in determining the intensity and risk of disease transmission [16]. Also, in our model we use transmission rates that explicitly incorporate mosquito biting rates, probabilities of virus transmission, densities of both populations, and host availability [1].

We use baseline parameter sets for Dengue and Chikungunya obtained from the current literature to perform a sensitivity analysis on quantities of interest how the reproduction numbers and the *I*_2*c*_ class.

## 2 Material and Methods

Python scripts were written to simulate the model with parameters in Table 1 and the MatPlotLib graphics plotting library was used to produce the figures for the diseases prevalence.

**Table 1:** Parameters for the model. The parameters for Dengue have the subscript 1, the parameters for Chikungunya have the subscript 2, with baseline values or range, and references.

| Symbol | Parameter definition | Value |
| --- | --- | --- |
| $h$ | Host recruitment rate | variable |
| $\mu^{-1}$ | Host lifespan | 75 years [1] |
| $\sigma_{vi}$ | Number of bites that one mosquito would give a human per unit time | 0.33-1.0 [1] |
| $\sigma_{hi}$ | The maximum number of mosquito bites that a human can suffer per unit time | 0.1-50 [1] |
| $\beta_{hvi}$ | Probability of virus transmission from an infectious vector to a susceptible human due to contact between both | 0.001-0.9 [1] |
| $\beta_{vhi}$ | Probability of virus transmission from an infectious human to a susceptible vector due to contact between both | 0.1-0.9 [1] |
| $\gamma_1^{-1}$ | Mean length of infectious period of Dengue in host | 10 days[23] |
| $\gamma_2^{-1}$ | Mean length of infectious period of Chikungunya in host | 7 days [4] |
| $\gamma_3^{-1}$ | Fraction of recovered individuals after acute phase of Chikungunya | $0.5\gamma_2^{-1}$ [41] |
| $\gamma_4^{-1}$ | Fraction of sub-acute infected individuals of Chikungunya | $0.5\gamma_2^{-1}$ [41] |
| $q$ | Vector recruitment rate | variable |
| $\eta_1^{-1}$ | Mean length of exposed period of Dengue in host | 4 days[24] |
| $\eta_2^{-1}$ | Mean length of exposed period of Chikungunya virus in host | 4 days[25] |
| $\delta^{-1}$ | Vector lifespan | 13 days[1] |
| $\sigma_1$ | Induced susceptibility to class $I_{2c}$ and $I_{3c}$ | 0-10 [2] |
| $\sigma_2$ | Induced susceptibility to class $R_c$ | 0-10 [2] |
| $\sigma_3$ | Induced susceptibility to class $R_d$ | 0-10 [2] |
| $\nu^{-1}$ | Mean length of sub-acute phase | 90 days [40] |

R scripts were written to process the uncertain parameters through Latin Hypercube Sampling algorithm from uniform and triangular probability distributions, 3000 samples were taken for produce each histogram and 5000 samples were taken for get the partial rank correlation coefficients (PRCCs).

We collect Dengue and Chikungunya Mexico 2014, 2015 and 2016 data from the registers in [14].

## 3 Calculation

In this section we present the formulation of a model that describes the transmission dynamics of Dengue and Chikungunya viruses where only a single vector, *Ae. aegypti*, is active. The population of hosts and vectors is assumed variable, although asymptotically constant, we also incorporate an exposed class in the humans as well as the possibility of re-infections by the second virus in those individuals already recovered from the first disease or those susceptible because clinical deterioration.

The population of human hosts is divided in the following classes: susceptible (*S*), virus 1 exposed (Dengue) (*E_d_*) (infected, but not infectious), virus 2 exposed (Chikungunya) (*E_c_*), virus 1 infected (*I_d_*), virus 2 infected (*I_c_*), recovered from virus 1 (*R_d_*), recovered from virus 2 (*R_c_*), virus 2 infected in sub-acute phase (*I*_2_*_c_*), and virus 2 infected in chronic phase (*I*_3_*_c_*); all this categories represent the first infections. To describe secondary infections by a different virus we add the next categories: virus (*I*_c_*_d_*).virus 2 second infected (*I_dc_*) virus 2 second infected in sub-acute phase (*I_d_*_2_*_c_*), virus 2 second infected in chronic phase (*I_d_*_3_*_c_*), recovered from both viruses (*R_dc_*) and recovered from virus 1 that he/she had or still he/she has Chikugunya (*R_cd_*).

Humans are born with a per-capita rate *h* and enter into the susceptible class, *S*. After an infectious mosquito bites a susceptible human who becomes infected, the individuals pass from the susceptible class *S* to the exposed class, *E_d_* or *E_c_*, respectively. After an intrinsic incubation period (*η_i_*, i=1,2), they go forward to the infectious class, *I_d_* or *I_c_*, respectively, in these phases humans can infect susceptible mosquitoes when are bitten. Later, infectious humans are recovered and pass to the recovered class, *R_d_* or *R_c_* respectively. For Dengue, the recovered humans are immune for life to the specific serotype that they acquired, but only they have a brief period of immunity to the other three serotypes, this could take from 4 to 6 months [1]; in our model we consider that only a dominant Dengue serotype is transmitted, an scenario well documented in the Americas [17]. The per-capita natural death rate for the humans is *μ*.

In the case of Chikungunya, around 50% of the infected hosts are recovered after 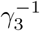 days and the other half enters to the sub-acute phase with 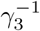 rate (since we suppose that 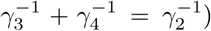, which lasts on average *ν*^−1^ days. Finally, they are moved into a chronic phase *I*_3_*_c_*. In these two last phases the patient can present clinical deterioration, making her/him more susceptible to other infections. Note that Chikungunya disease gives life-long immunity [18].

We allow that secondary infections take place only in the host and these are governed for the susceptibility coefficients *σ_i_, i* = 1, 2, 3; to explore this condition we add the last phases of each disease in our model.

Analogously, the fraction *q*(1 – *p_d_*) of the vector population (*Ae. aegypti*) that was born susceptible, is divided into the following classes: susceptible (*A*), infected with Dengue (*B_d_*) and infected with Chikungunya (*B_c_*), a susceptible mosquito becomes infected because it bites an infectious human of Dengue or Chikungunya, so that it pass to the infected class, *B_d_* or *B_c_*, respectively, the mosquito remains infectious for life (on average *δ*^−1^days). See de Fig. 1.

**Figure 1:**
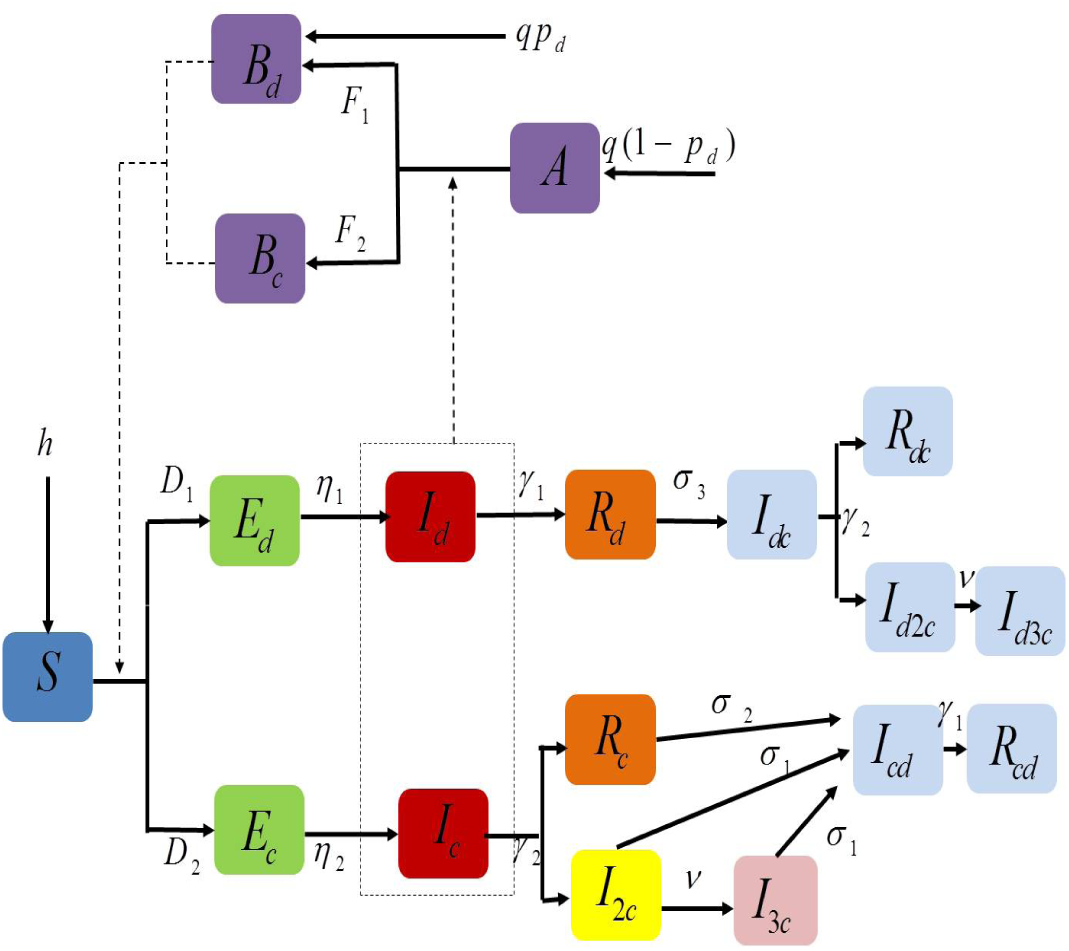
Graphic representation of the model. The black arrows indicate the diseases transitions and the dashed arrows indicate the contacts between humans and vectors. The human population is divided into susceptible (S), exposed of Dengue (*E_d_*), exposed of Chikungunya (*E_c_*), Dengue infected (*I_d_*),Chikungunya infected (*I_c_*), recovered from Dengue (*R_d_*), recovered from Chikungunya (*R_c_*), Chikungunya infected in sub-acute phase (*I*_2*c*_) and Chikungunya infected in chronic phase (*I*_3*c*_). The vector population is divided into susceptible (*A*), infected with Dengue (*B_d_*) and infected with Chikungunya (*B_c_*). The categories in light blue indicate compartments at role of secondary infections. A full description of the parameters can be found in the next section.

### 3.1 Model Equations

In this model the human host population resides in a region where *Ae. aegypti* is present and can transmit the two viruses. We have the following equations:

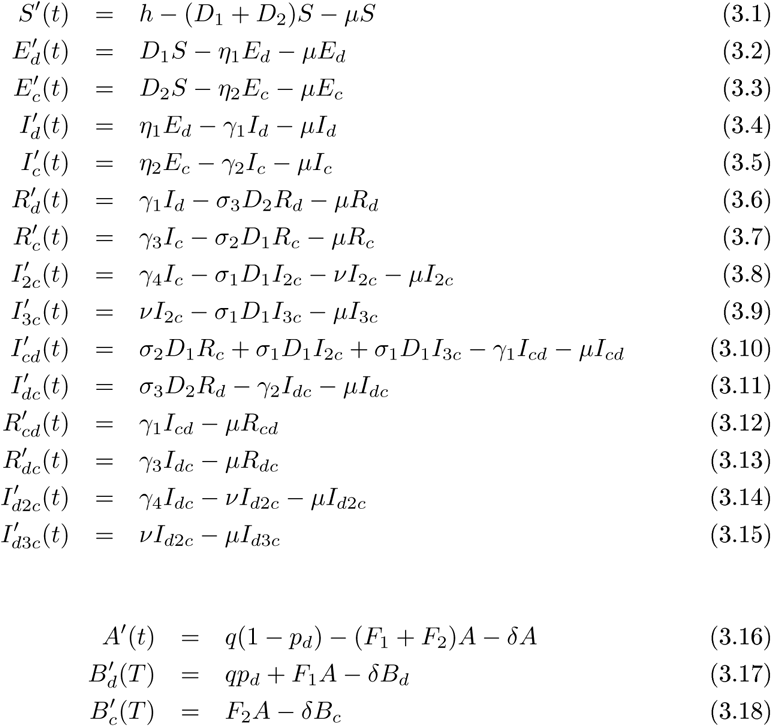
with *p_d_* because the vertical transmission of Dengue in the mosquito, where the parameters with subscript 1 refer to Dengue and those with subscript 2 refer to Chikungunya. Also, we have that *N* = *S* + *E_c_* + *E_d_* + *I_d_* + *I_c_* + *R_d_* +*R_c_* + *I*_2_*_c_* + *I*_3_*_c_* + *I_cd_* + *I_dc_* + *R_cd_* + *R_dc_* + *I_d_*_2_*_c_* + *I_d_*_3_*_c_* and *M* = *A* + *B_d_* + *B_c_* are the total population sizes for hosts and vectors respectively (see Table 1 for parameters definitions and values).

First human infections of the corresponding viruses are carried out at rates

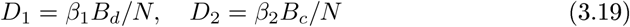
(vector to host transmission). First infections in vectors are carried out at rates

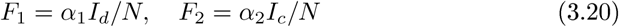

Where

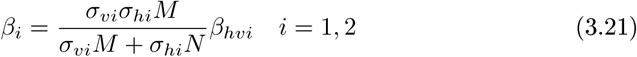
and

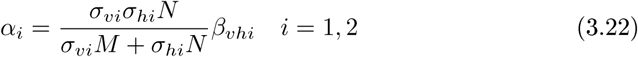

We use these contact rates that follow the hypothesis of Manore et al. [1], where *σ_υi_* is the maximum number of bites that one mosquito would give a human per unit time (related to time of egg production), and *σ_hi_* is the maximum number of bites that a human can suffer per unit time. Then, *σ_υi_M* is the optimal number of bites that a mosquito searches per unit time and *σ_hi_N* is the maximum available number of bites in humans per unit time [1].

The total number of vector-host contacts depends of both population densities [1] and is defined how:

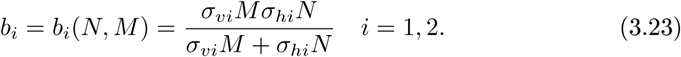

Also we define *b_hi_* = *b_hi_*(*N*,*M*) = *b_i_*(*N*,*M*)/*N* as the number of bites that a human suffers per unit time, and *b_υi_* = *b_υi_*(*N*,*M*) = *b_i_*(*N*,*M*)*/M* as the number of bites that a mosquito gives per unit time [1]. With this kind of biting rate we can manage the entire spectrum of possible vector-to-host ratios in contrast with standard frequency-dependent contact rates [1].

We define *D_i_, i* = 1, 2, how the force of infection from mosquitoes to humans, which is the product of the number of bites that a human suffers by mosquitoes per unit time, *b_hi_*, the probability that the mosquito is infectious, *B_d_*/*N* or *B_c_*/*N*, respectively, and the probability of virus transmission from an infected mosquito to a susceptible human, *β_hυi_* [1]. Now, *F_i_, i* = 1, 2, is the force of infection from humans to mosquitoes, which is defined as the number of bites that a mosquito gives to a human per unit time, *b_υi_*, the probability that the human is infectious, *I_d_*/*N* or *I_c_*/*N*, respectively, and the probability of virus transmission from an infected human to a susceptible mosquito, *β_υhi_* [1].

## 4 Results

Python scripts were written to simulate the model with parameters in Table 1 and the MatPlotLib graphics plotting library was used to produce the figures for the diseases prevalence.

R scripts were written to process the uncertain parameters through Latin Hypercube Sampling algorithm from uniform and triangular probability distributions, 3000 samples were taken for produce each histogram and 5000 samples were taken for get the partial rank correlation coefficients (PRCCs)

### 4.1 The basic reproduction number

Note that the equation for the total host population is

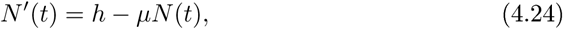
we have that as *t* → ∞ then *N*(*t*) → *h*/*μ*.

Moreover, the equation for the total vector population is

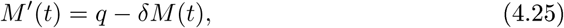
so we have that as *t* → ∞, then *M*(*t*) → *q*/*δ*. Hence we can substitute *A* = *q*/*δ* – *B_d_* – *B_c_* and we reduce the system (3.1)-(3.18).

We consider the set

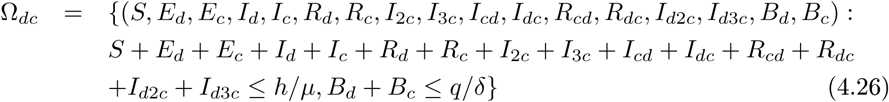
which is bounded by the total host and vector population, when the diseases are not present.

The basic reproduction number is the total number of secondary infections caused by the introduction of an infected individual in a population fully susceptible, this threshold parameter helps to predict the spread or extinction of the virus within the susceptible population [2].

To calculate the basic reproduction number for our model, we proceed as in [2], we obtain the next-generation matrix, then it is evaluated at the disease-free equilibrium given by *D*_1_ = *D*_2_ = *F*_1_ = *F*_2_ = 0; in our case we assume *p_d_* = 0, thus:

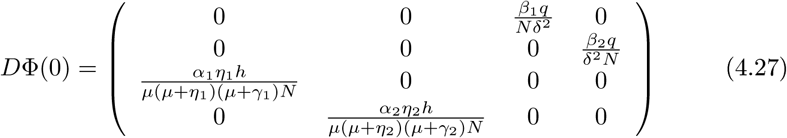

The basic reproduction number is therefore

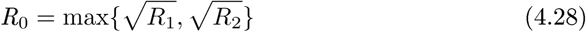
with

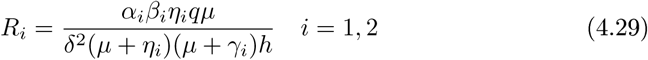

We can interpret this quantity as follows: the mosquitoes acquire the infection by the virus 1 (Dengue) at the infection rate 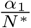 after they take a blood meal from an infected individual. A Dengue infected individual generate 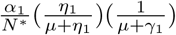 vector infections; this is the product between the Dengue infection rate for infected individuals, the average duration in Dengue exposed class and the average duration in the Dengue infected class. So, Dengue infected individuals generate the next average number of new vector infections (note that *A*\* = *q*/*δ* and *N*\* = *h*/*μ*):

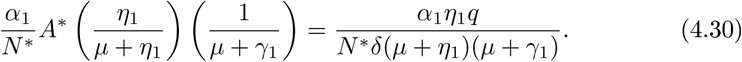

Similarly, the virus 2 (Chikungunya) infected individuals generate the next average number of new vector infections:

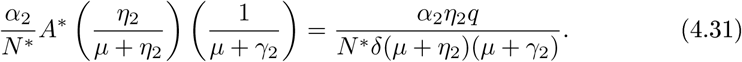

The individuals acquire the infection by the virus 1 (Dengue) at the infection rate 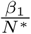 after they are bitten by an infected vector. A Dengue infected vector generate 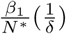 humans infections; this is the product between the Dengue infection rate for infected vectors and their average duration in the Dengue infected class. So, Dengue infected vectors generate the next average number of new humans infections (note that *S*\* = *h*/*μ*):

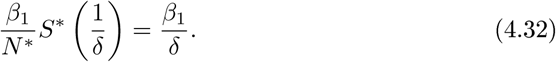

Similarly, the virus 2 (Chikungunya) infected vectors generate the next average number of new humans infections:

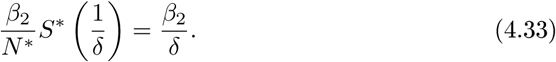

Observe that the geometric mean of (4.30)-(4.32) gives the virus 1 (Dengue) basic reproduction number and the geometric mean of (4.31)-(4.33) gives the virus 2 (Chikungunya) basic reproduction number; so, the basic reproduction number for our model is the maximum of the reproduction numbers of each virus.

Since the basic reproduction number is a threshold parameter for our model, then if *R*_0_ > 1, the disease may spread into the host population. Otherwise, if *R*_0_ ≤ 1 the disease will die out from the host population [2].

### 4.2 Numerical Simulations

This section presents the numerical simulations of the model dynamics, as well as a sensitivity analysis of basic reproduction numbers of both diseases.

We performed numerical simulations to explore the prevalence of Dengue and Chikungunya virus using the parameters shown in Table 1 and too we used them as a baseline for the sensitivity analysis. For the simulations and the uncertainty and sensitivity analysis we used: *h* = 2.55, *q* = 8076.92 since *N* = *h*/*μ* and *M* = *q*/*δ*; which means that N is around 70 000 individuals, also we set that the mosquito population is 1.5 times the host population; moreover, *p_d_* = 9.81*e* – 3, because the fraction of mosquitoes that was born infected is small and this is the result from our parameters estimation for Mexico (see Table 2). We assume *σ*_1_ > *σ*_2_ because a larger susceptibility to other infections occurs by clinical deterioration in the classes *I*_2_*_c_* and *I*_3_*_c_*. Moreover, we suppose that 50% of Chikungunya infected are recovered (*γ*_3_ in our model) and the other half (*γ*_4_ in our model) passes to the sub-acute phase [41]. Both populations were normalized to 1 and we used as initial conditions: *S* = 0.98, *I_d_* = 0.01 *I_c_* = 0.01 *A* = 1, and the other classes equals to zero, this because we assumed there is already a small fraction of infected with each virus in the population.

**Table 2:** Estimated Parameters. Subscript 1 for Dengue parameters and subscript 2 for Chikungunya parameters.

| Parameter | Value | Parameter | Value |
| --- | --- | --- | --- |
| $\sigma_{v1}$ | 5.88e-01 | $\sigma_{v2}$ | 8.41e-01 |
| $\sigma_{h1}$ | 2.05e01 | $\sigma_{h2}$ | 2.07e1 |
| $\beta_{hv1}$ | 3.05e-02 | $\beta_{hv2}$ | 4.34e-02 |
| $\beta_{vh1}$ | 3.84e-01 | $\beta_{vh2}$ | 3.58e-01 |
| $\gamma_1$ | 8.33e-02 | $\gamma_2$ | 1.43e-01 |
| $\eta_1$ | 2.44e-01 | $\eta_2$ | 3.33e-01 |
| $h$ | 8.98e-01 | $\mu$ | 3.42e-05 |
| $\delta$ | 5.0e-02 | $p_d$ | 9.81e-03 |

We want to analyze the effect of change the basic reproduction numbers on diseases prevalence, we took infection rates values into the baseline of the parameters (see Table 1) and then we compared in size *R*_1_, Dengue basic reproduction number with *R*_2_, Chikungunya basic reproduction number.

Fig. 2 shows the total number of infected individuals for each virus *I_d_* and *I_c_* For a period of 60 days, here we assume (a)*p_d_* = 0 and (b)*p_d_* = 9.81*e* – 3. Both infected populations are shown for the case where the Chikungunya basic reproduction number is larger than the Dengue basic reproduction number (*R*_1_ < *R*_2_), with *R*_1_ = 2.95, *R*_2_ = 4.14, so we will have *β_hυ_*_1_ = 0.4, *β_υh_*_1_ = 0.32, *β_hυ_*_2_ = *β_υh_*_2_ = 0.6, *σ_υi_* = 0.5, and *σ_hi_* = 19, *i* = 1, 2, note that all parameters values are in the range of the baseline parameters (Table 1). Observe that Chikungunya prevalence is larger than Dengue’s for all the period and the epidemic peak occurs around the same time for both diseases.

**Figure 2:**
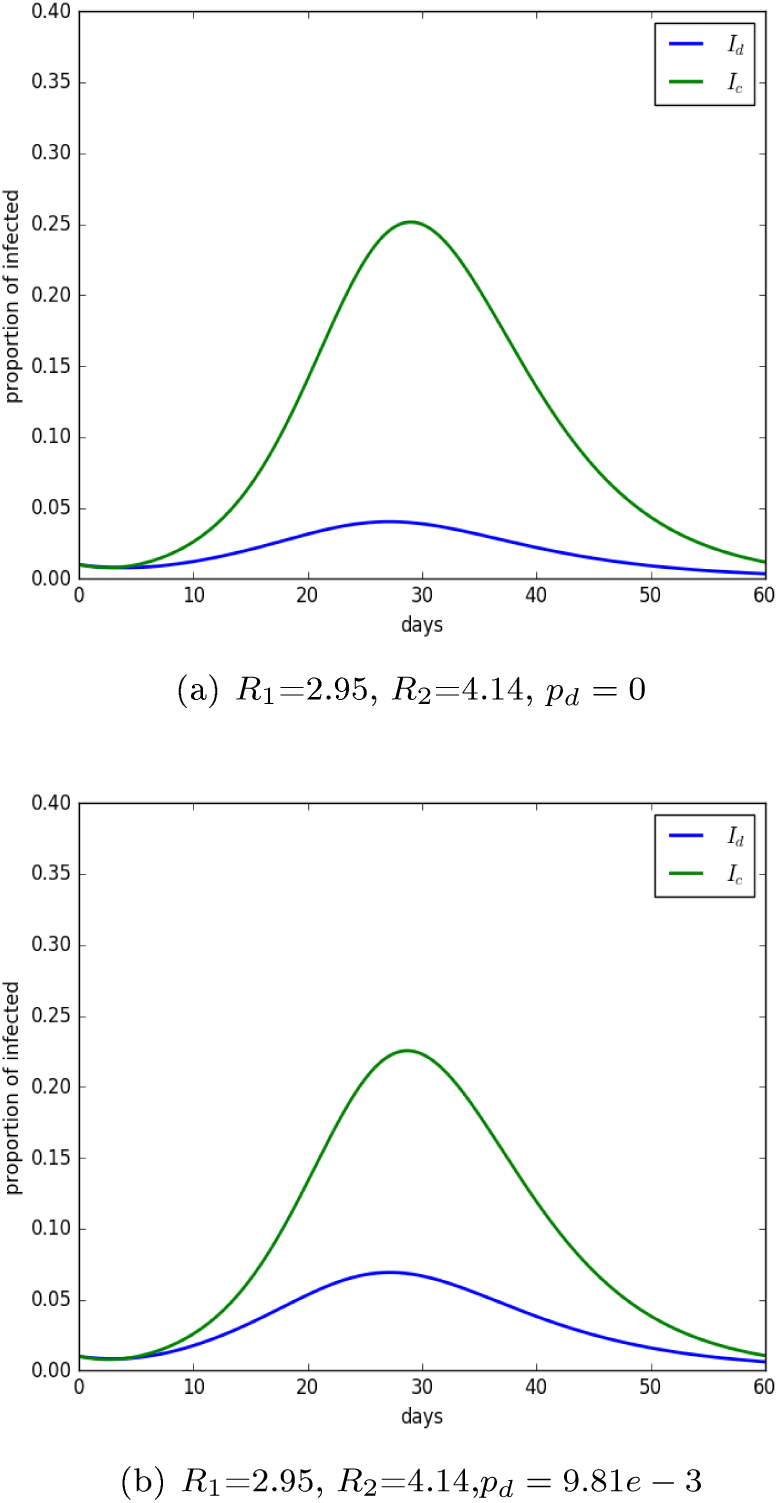
Disease progression of Dengue and Chikungunya. The *y*-axis is the proportion of hots and the *x*-axis is time in days. Baseline parameters from Table 1 are used with initial conditions (*S*,*E_d_*,*E_c_*,*I_d_, I_c_, R_d_, R_c_, I*_2*c*_, *I*_3*c*_, *I_cd_, I_dc_, R_cd_, R_dc_, I_d_*_2_*_c_, I_d_*_3_*_c_, A, B_d_, B_c_*) = (0.98, 0, 0, 0.01, 0.01, 0, 0, 0, 0, 0, 0, 0, 0, 0, 0, 1, 0, 0); moreover, we assume various *R_i_* values, *i* = 1, 2, with *σ*_1_ = 2, *σ*_2_ = 1 and *σ*_3_ = 1.

Fig. 3 we show *I_d_* and *I*_c_ for a period of 80 days. In this case the basic reproduction number of Chikungunya is less than the one of Dengue (*R*_2_<*R*_1_), so we take *R*_1_=3.71 and *R*_2_=2.76, so we will have *β_hυ_*_1_ = *β_υh_*_1_ = 0.45, *β_hυ_*_2_ = *β_υh_*_1_ = 0.4, with *σ_υi_* = 0.5 and *σ_hi_* = 19, *i* = 1, 2, note that all this parameters values are in the range of baseline parameters (Table 1). In this case Chikungunya prevalence is lower than the Dengue prevalence and the Dengue epidemic peak occurs after Chikungunya peak.

**Figure 3:**
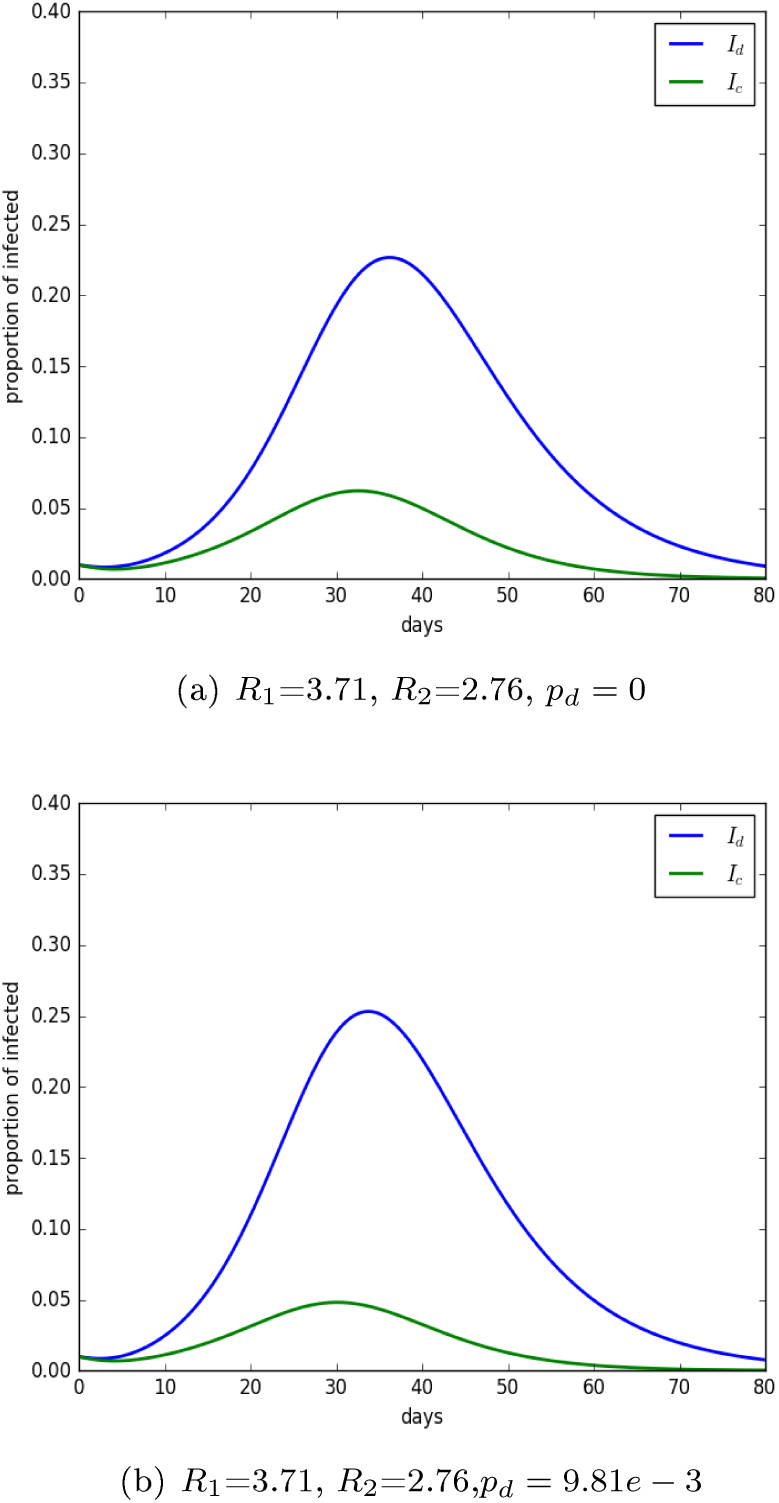
Disease progression of Dengue and Chikungunya. The *y*-axis is the proportion of hots and the *x*-axis is time in days. Baseline parameters from Table 1 are used with initial conditions (*S, E_d_, E_c_, I_d_, I_c_, R_d_, R_c_, I*_2_*_c_, I*_3_*_c_, I_cd_, I_dc_, R_cd_, R_dc_, I_d_*_2_*_c_, I_d_*_3_*_c_, A, B_d_, B_c_*) = (0.98, 0, 0, 0.01, 0.01,0, 0, 0, 0, 0, 0, 0, 0, 0, 0, 1, 0, 0): moreover, we assume various *R_i_* values, *i* = 1, 2, with *σ*_1_ = 2, *σ*_2_ = 1 and *σ*_3_ = 1.

So far we observe the importance of consider vertical transmission since in the previous figures it is shown that although the parameter *p_d_* take small values, it has an impact on the Dengue prevalence, more precisely, *p_d_* increases it. Recent studies in Mexico report a vertical transmission around of 1.2%, which is very similar to the correspond obtained by us of *p_d_* = 9.81*e* – 3 it due to our model fit to data (see Table 2).

On the other hand we too analyze what parameters involved in infection forces have more influence on reduce susceptible host population, turns out that these parameters are: *σ*_*υ*1_ and *β_hυ_*_1_, i-e. the number of times of one mosquito would bites a human per unit time and the probability of virus transmission from infected vectors to susceptible humans, respectively.

Moreover, we compare our model results with data from Mexico 2015 [14]

Fig. 4 shows the total number of infected individuals for each virus; Dengue and Chikungunya in a period of 52 weeks. We compared the predictions of our model for a period of 34 weeks, namely, the last outbreak of the year; we found a parameters set that allow our model fits the given data, for this we used a nonlinear least-squares procedure as in [19], and we minimized the sum of the squared errors (*SEDC*) given for the next function.

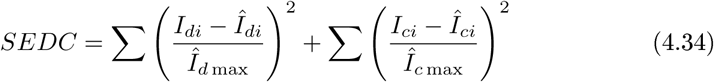

**Figure 4:**
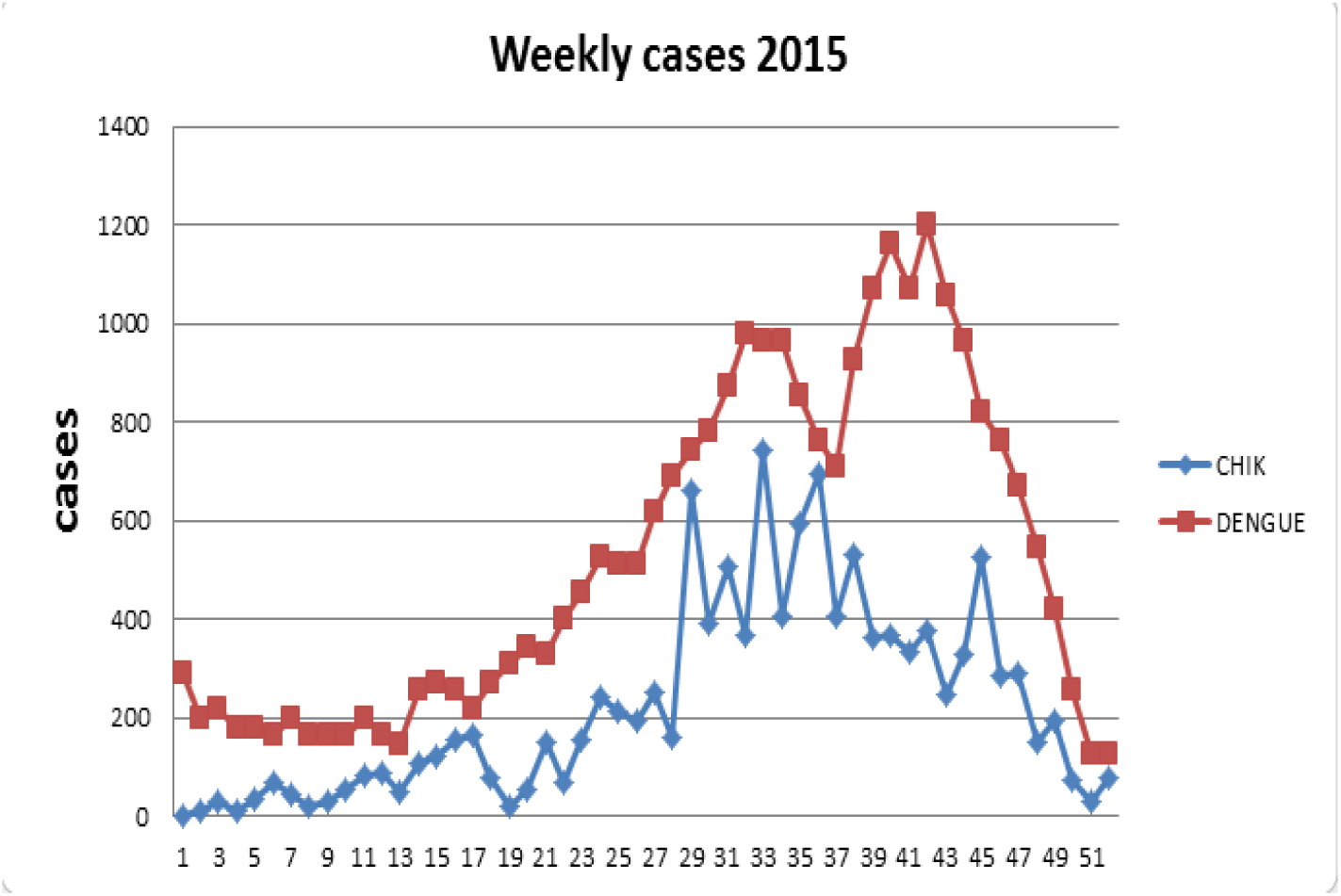
Weekly cases of Dengue and Chikungunya in Mexico 2015 [14]. The *y*-axis is the number of infected individuals and the *x*-axis is time in weeks.

Where 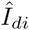 are the data of Dengue infected in Mexico, 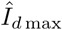 is the maximum number of infected during the year and *I_di_* are the results from the model, analogously, we used these definitions for Chikungunya infected.

We obtained the next parameters set (Table 2) that make the model fits to the data from Mexico of Dengue and Chikungunya infected.

Fig. 6 shows the total number of infected individuals for each virus *I_d_* and *I_c_* for a period of 370 days, we observe that the model fits well to the data with the parameters in Table 2; with these parameters we have that the Dengue basic reproduction number is *R*_1_=1.42, and the Chikungunya basic reproduction number is *R*_2_=1.75, these amounts are given by the fitted model.

**Figure 5:**
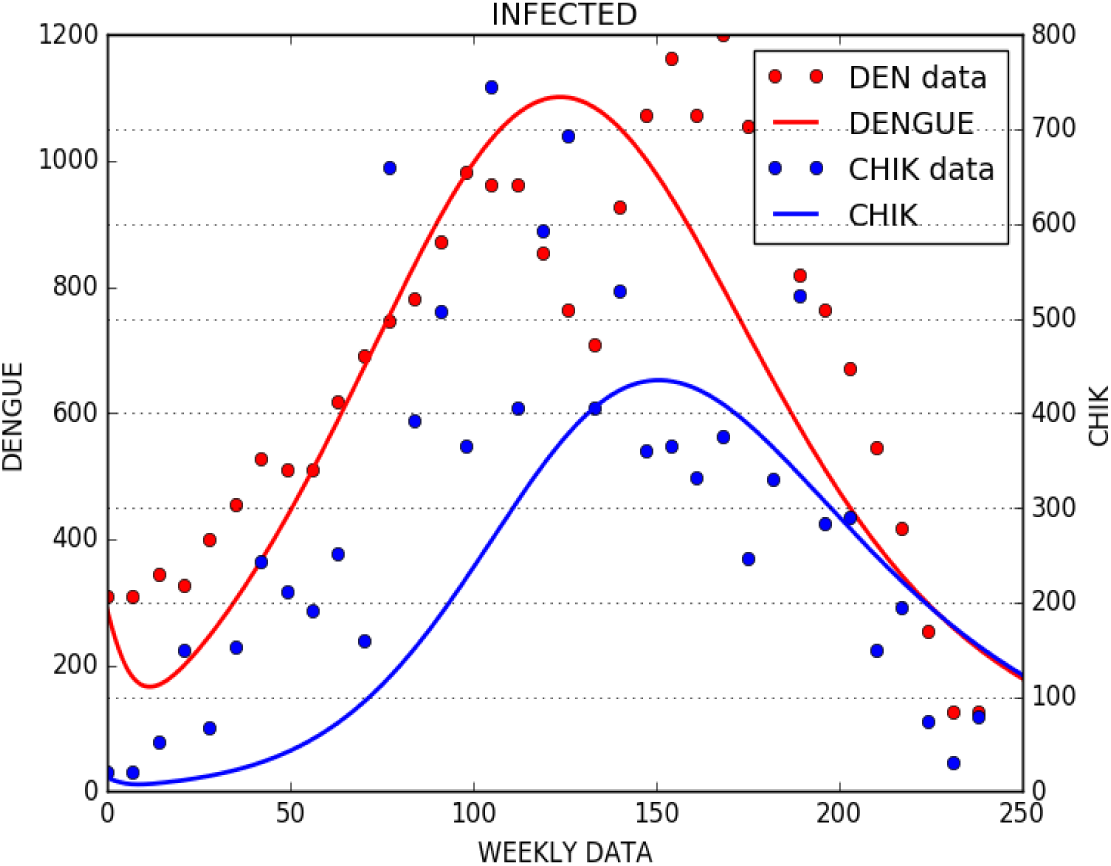
Weekly cases of Dengue and Chikungunya in Mexico 2015 [14] along with the model predictions. The *y*-axis is the number of infected individuals and the *x*-axis is time in days. The red points are the Dengue data, the blue ones are the Chikungunya data, and the corresponding lines are the results of our model when the *SEDC* function is minimized.

**Figure6:**
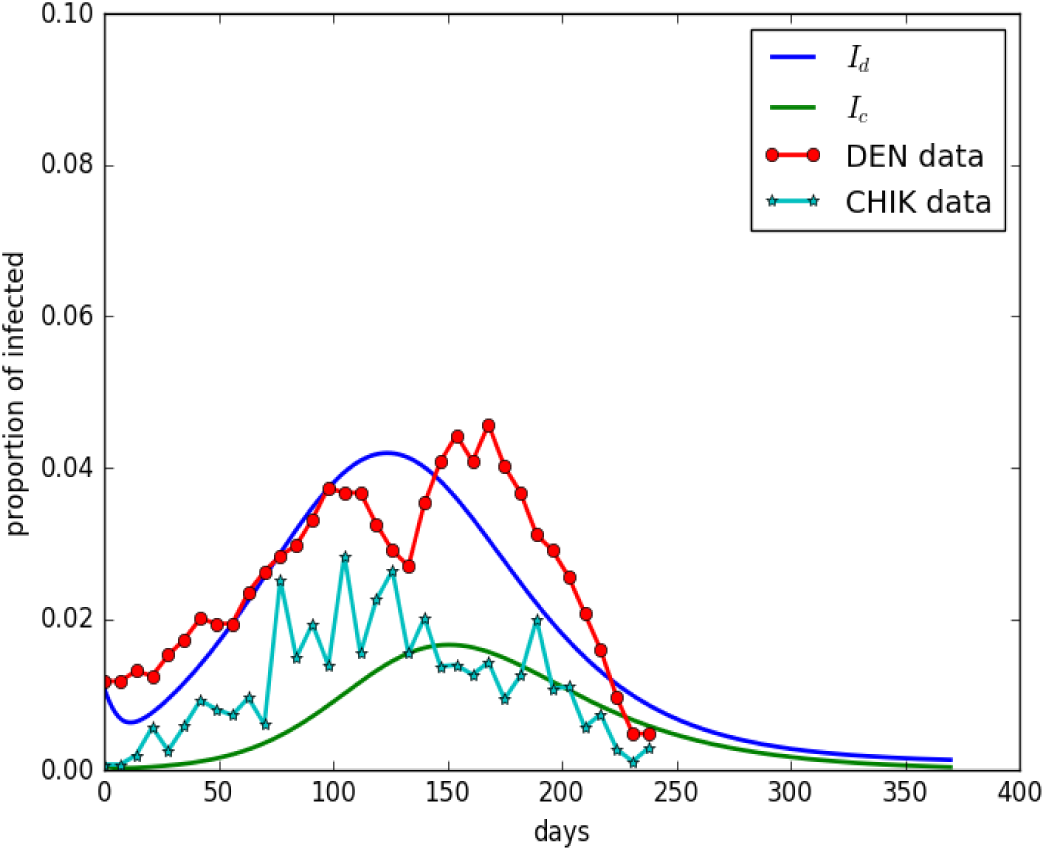
Disease progression of Dengue and Chikungunya along with weekly cases in Mexico 2015 [14]. The *y*-axis is the number of infected individuals and the *x*-axis is time in days. Baseline parameters from Table 2 are used with initial conditions (*S, E_d_, E_c_, I_d_, I_c_, R_d_, R_c_, I*_2_*_c_, I*_3_*_c_, I_cd_, I_dc_, R_cd_, R_dc_, I_d_*_2_*_c_, I_d_*_3_*_c_, A, B_d_, B_c_*) = (1 – 305(*μ*/*h*), 0, 0, 290(*μ/h*)*, 15*(*μ/h*), 0, 0, 0, 0, 0, 0, 0, 0, 0, 0, 1, 0, 0), with *σ*_1_=2, *σ*_2_= 1 and *σ*_3_=1.

Note in the Fig. 6 that Dengue data behavior seems anomalous because it shows two peaks at the end of the year, besides that the last peak decays very fast in the time, this may be one reason why our model does not fit the last few weeks of Dengue data; however this behavior is observed in other years in México (2013, 2010 [14]), therefore we decide to adjust our model only for Dengue data from 2013 (the lasts 24 weeks), since in this year there was not Chikungunya disease in the country; we obtain a parameters set for our model that fits these data.

In Table 3 we give the parameters set that makes the model fits to the data from Mexico 2013 of Dengue infected.

**Table 3:**
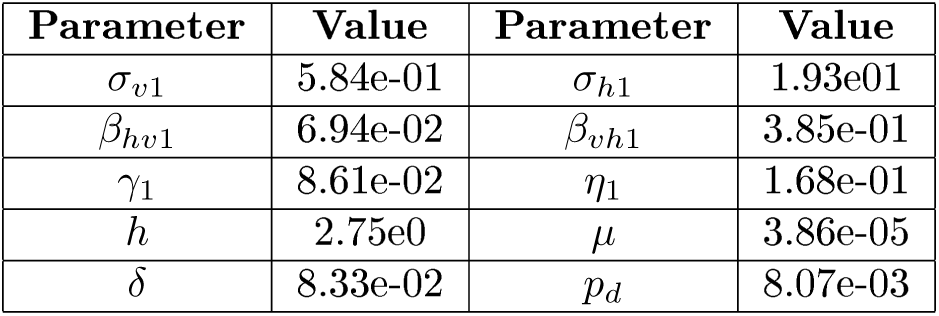
Estimated Parameters. Subscript 1 for Dengue parameters.

Accord the results in Fig. 6 and 7 we conclude that the presence of Chikungunya disease does not have influence over Dengue behavior, since we see the same trend for this disease in some other years; we are looking for the cause of this trend, we thought that maybe it is due to extreme mobility factors or climatology conditions.

**Figure7:**
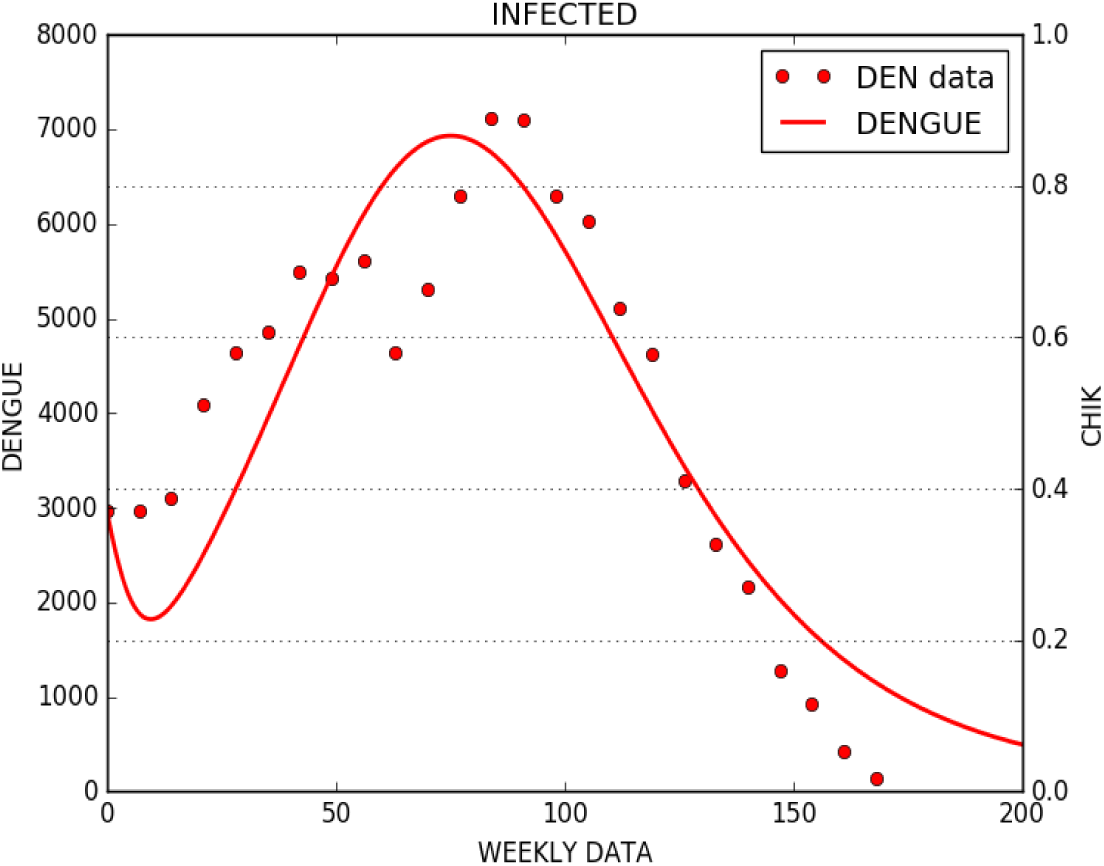
Weekly cases of Dengue in Mexico 2013 [14] along with the model predictions. The *y*-axis is the number of infected individuals and the *x*-axis is time in days. The red points are the Dengue data and the corresponding lines are the results of our model when the *SEDC* function is minimized.

On the other hand, we decide to analyze the outputs from our model with box plots in order to visualize the behavior of the predictions in some specifics times. We perform a sampling based on Latin Hypercube [22] to generate initial conditions for both diseases (only for the categories *I_d_* and *I_c_*), we suppose that each initial condition follows a uniform distribution in a given range (See Table 4); moreover, we use the parameters set in Table 2.

**Table 4:**
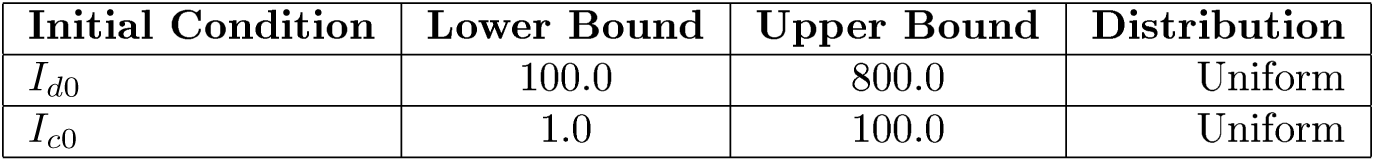
Parameter range for sampling initial conditions.

**Table 5:** Statistics of model predictions for Dengue. The sample times are time 1=30, time 2=60, time 3=90, time 4=120, time 5=150, time 6=180 and time 7=238 all in days.

|  | TIME 1 | TIME 2 | TIME 3 | TIME 4 | TIME 5 | TIME 6 | TIME 7 |
| --- | --- | --- | --- | --- | --- | --- | --- |
| Lower whisker | 133.11 | 328.99 | 515.07 | 490.91 | 303.34 | 152.46 | 44.94 |
| Lower hinge | 243.7 | 514.71 | 787.75 | 803.13 | 536.42 | 291.54 | 84.66 |
| Median | 348.92 | 664.56 | 938.4 | 930.1 | 666.24 | 383.56 | 114.61 |
| Upper hinge | 451.17 | 795.83 | 1055.5 | 1082.95 | 863.62 | 549.09 | 179.57 |
| Upper whisker | 552.04 | 944.52 | 1277.86 | 1356.16 | 1301.88 | 935.39 | 321.49 |

**Table 6:** Statistics of model predictions for Chikungunya. The sample times are time 1=30, time 2=60, time 3=90, time 4=120, time 5=150, time 6=180 and time 7=238 all in days.

|  | TIME 1 | TIME 2 | TIME 3 | TIME 4 | TIME 5 | TIME 6 | TIME 7 |
| --- | --- | --- | --- | --- | --- | --- | --- |
| Lower whisker | 1.16 | 3.57 | 8.19 | 13.45 | 16.38 | 91.33 | 44.58 |
| Lower hinge | 28.49 | 96.39 | 236.86 | 381.67 | 390.15 | 272.66 | 88.78 |
| Median | 55.44 | 181.91 | 416.94 | 581.63 | 508.06 | 328.78 | 98.99 |
| Upper hinge | 82.04 | 262.73 | 570.02 | 766.44 | 648.6 | 394.08 | 118.3 |
| Upper whisker | 114.97 | 404.98 | 957.23 | 1180.69 | 879.58 | 572.73 | 162.27 |

We generated 3000 samples to produce the following results.

In the Fig. 8 and 9 it is shown that the data of Chikungunya and Dengue from Mexico 2015 are between the outputs from our model, this fits much better to Chikungunya data and the same happens with the first peak of Dengue epidemic, we will continue searching which is the origin of the apparent second peak that it is appreciated in the Dengue data, which our model can not fit. In general, we can predict the fate for Dengue and Chikungunya with our work.

**Figure 8:**
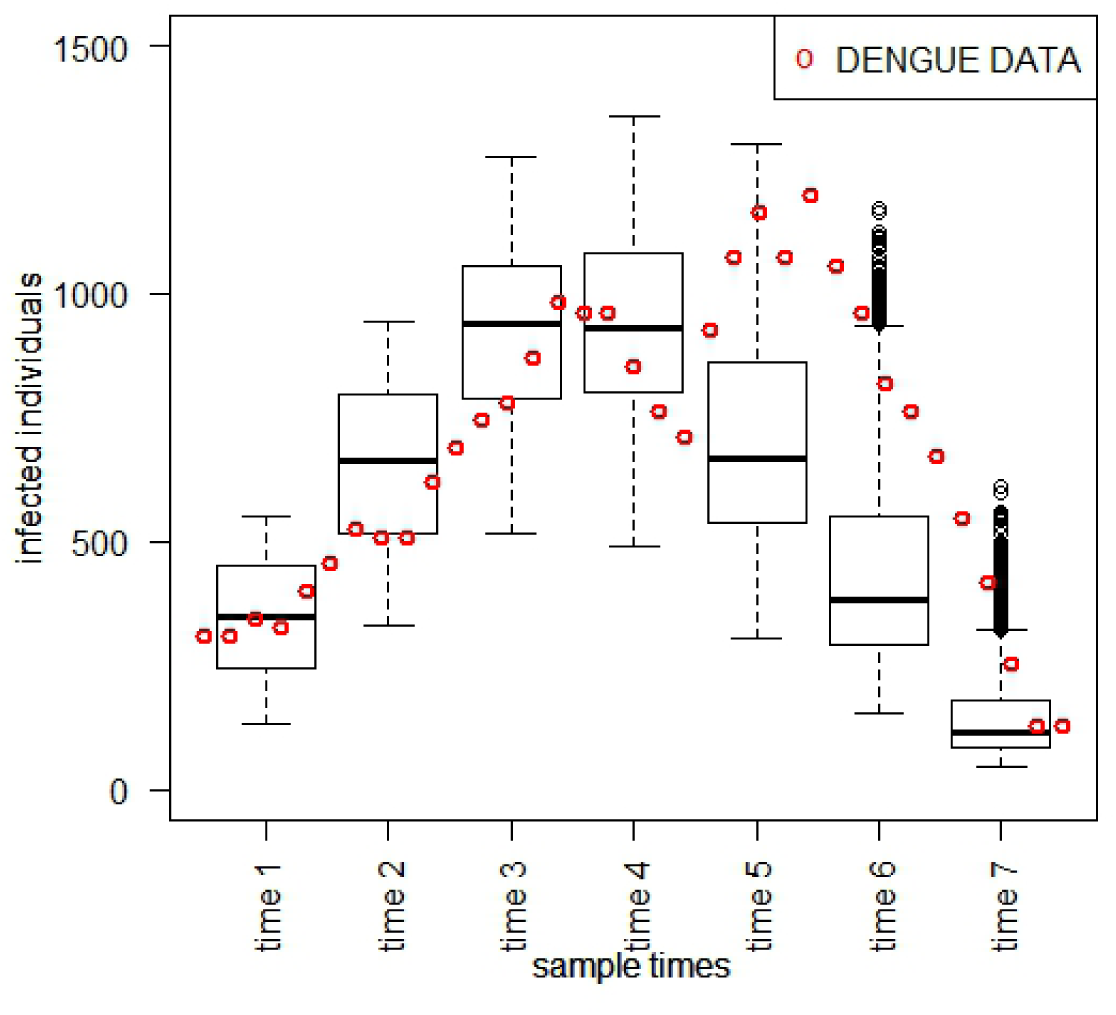
Weekly cases of Dengue in Mexico 2015 [14] along with the box plots of model predictions at seven different times for 3000 samples. The *y*-axis is the number of infected individuals and the *x*-axis is time in days. The red points are the Dengue data and the box plots are in time 1=30, time 2=60, time 3=90, time 4=120, time 5=150, time 6=180 and time 7=238.

**Figure 9:**
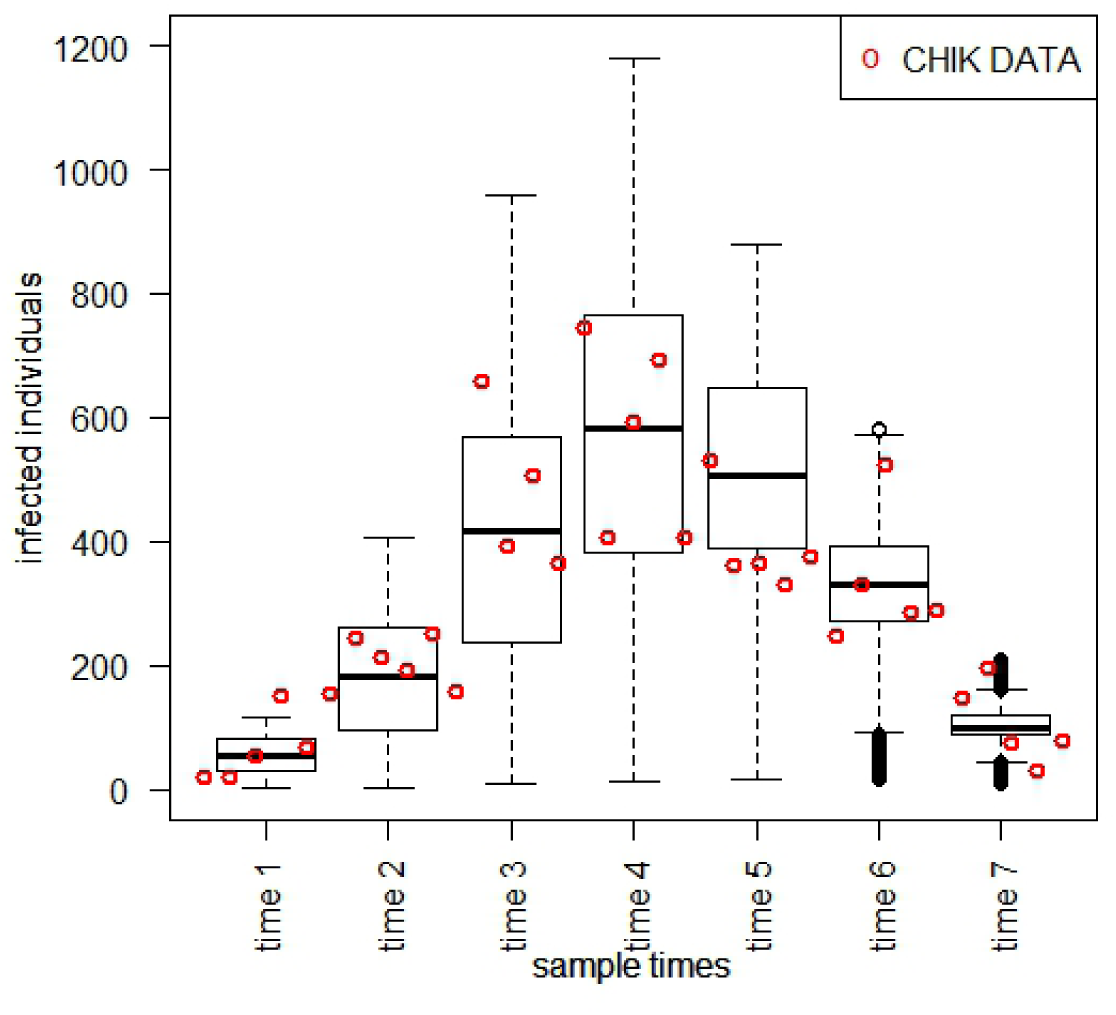
Weekly cases of Chikungunya in Mexico 2015 [14] along with the box plots of model predictions at seven different times for 3000 samples. The *y*-axis is the number of infected individuals and the *x*-axis is time in days. The red points are the Chikungunya data and the box plots are in time 1=30, time 2=60, time 3=90, time 4=120, time 5=150, time 6=180 and time 7=238.

### 4.3 Sensitivity Analysis

In this section we present an analysis to investigate the sensitivity of both basic reproduction numbers to our choice of model parameters. We chose a probability density function (pdf) for each parameter, after, we give a specific range of possible values that the pdf can take with some degree of probability [21]. We used a triangular distribution function as it takes into account that the most likely values are near of its peak, and the uniform distribution when the probability of occurrence is even for any interval with the same length into the uniform distribution support [21]. We used a method based on Latin Hypercube Sampling [22] to generate the model parameters. In our case, we chose either a uniform or a triangular pdf for each parameter (see Tables 7 and 8), moreover we assume that the parameter 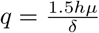, since *M* = 1.5*N*.

**Table 7:** Parameter range for sampling *R*_1_. Symmetric Triangular distributions have the average value at the fixed point shown in Table 1.

| Parameter | Lower Bound | Upper Bound | Distribution |
| --- | --- | --- | --- |
| $\sigma_{v1}$ [1] | 0.33 | 1.0 | Uniform |
| $\sigma_{h1}$ [1] | 0.1 | 50 | Uniform |
| $\beta_{hv1}$ [1] | 0.001 | 0.9 | Uniform |
| $\beta_{vh1}$ [1] | 0.1 | 0.9 | Uniform |
| $h$ | 0.01 | 1.0 | Uniform |
| $\gamma_1$ [23] | 1.0/12.0 | 1.0/5.0 | Triangular |
| $\eta_1$ [24] | 1.0/8.0 | 1.0/3.0 | Triangular |
| $\mu$ [1] | 1.0/(80.0*365.0) | 1.0/(65.0*365.0) | Triangular |
| $\delta$ [1] | 1.0/20.0 | 1.0/12.0 | Triangular |

**Table 8:** Parameter range for sampling *R*_2_. Symmetric Triangular distributions have the average value at the fixed point shown in Table 1.

| Parameter | Lower Bound | Upper Bound | Distribution |
| --- | --- | --- | --- |
| $\sigma_{v2}$ [1] | 0.33 | 1.0 | Uniform |
| $\sigma_{h2}$ [1] | 0.1 | 50 | Uniform |
| $\beta_{hv2}$ [1] | 0.001 | 0.9 | Uniform |
| $\beta_{vh2}$ [1] | 0.1 | 0.9 | Uniform |
| $h$ | 0.01 | 1.0 | Uniform |
| $\gamma_2$ [4] | 1.0/9.0 | 1.0/2.0 | Triangular |
| $\eta_2$ [25] | 1.0/8.0 | 1.0/3.0 | Triangular |
| $\mu$ [1] | 1.0/(80.0*365.0) | 1.0/(65.0*365.0) | Triangular |
| $\delta$ [1] | 1.0/20.0 | 1.0/12.0 | Triangular |

**Table 9:**
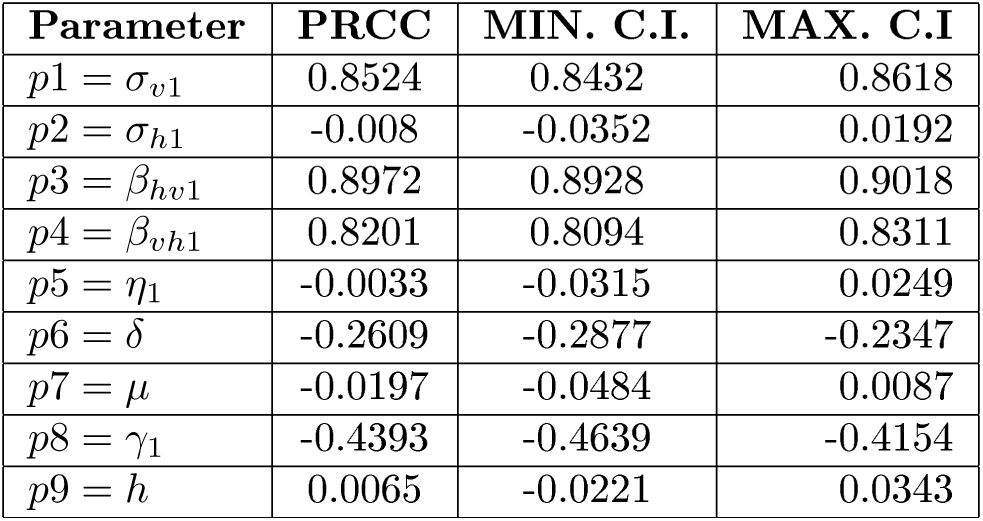
PRCCs and Confidence Intervals for Dengue basic reproduction number. These were calculated using Latin Hypercube Sampling method in each parameter ranges for the basic reproduction number of Dengue (see Fig. 11(a))

**Table 10:** PRCCs and Confidence Intervals for Chikungunya basic reproduction number. These were calculated using Latin Hypercube Sampling method in each parameter ranges for the basic reproduction number of Chikungunya (see Fig. 11(b))

| Parameter | PRCC | MIN. C.I. | MAX. C.I |
| --- | --- | --- | --- |
| $p1 = \sigma_{v2}$ | 0.8371 | 0.8263 | 0.8482 |
| $p2 = \sigma_{h2}$ | 0.0295 | 0.0025 | 0.0565 |
| $p3 = \beta_{hv2}$ | 0.8905 | 0.8854 | 0.8956 |
| $p4 = \beta_{vh2}$ | 0.806 | 0.7926 | 0.8193 |
| $p5 = \eta_2$ | 0.0044 | -0.0234 | 0.032 |
| $p6 = \delta$ | -0.2556 | -0.2828 | -0.2293 |
| $p7 = \mu$ | -0.0149 | -0.0427 | 0.0116 |
| $p8 = \gamma_2$ | -0.6208 | -0.642 | -0.6001 |
| $p9 = h$ | 0.0017 | -0.0252 | 0.0295 |

We generated 3000 sets of parameters to produce the following results.

The support for the distributions and methodology is based on published work by [24, 4, 25, 1]. In particular Pandey et. al. [24] estimated 7 total parameters for the vector-host model by Bayesian MCMC using the cumulative DHF prevalence data where they used uniform distributions as priors for all the parameters. Also Manore et. al. [1] performed a global uncertainty analysis using uniform distributions for the parameters ranges.

In Fig. 10 we show the histograms of Dengue and Chikungunya basic reproduction numbers respectively, where the mean is shown by a red point on the lower axis. These reproduction numbers agree with those reported in the literature[20, 25].

**Figure 10:**
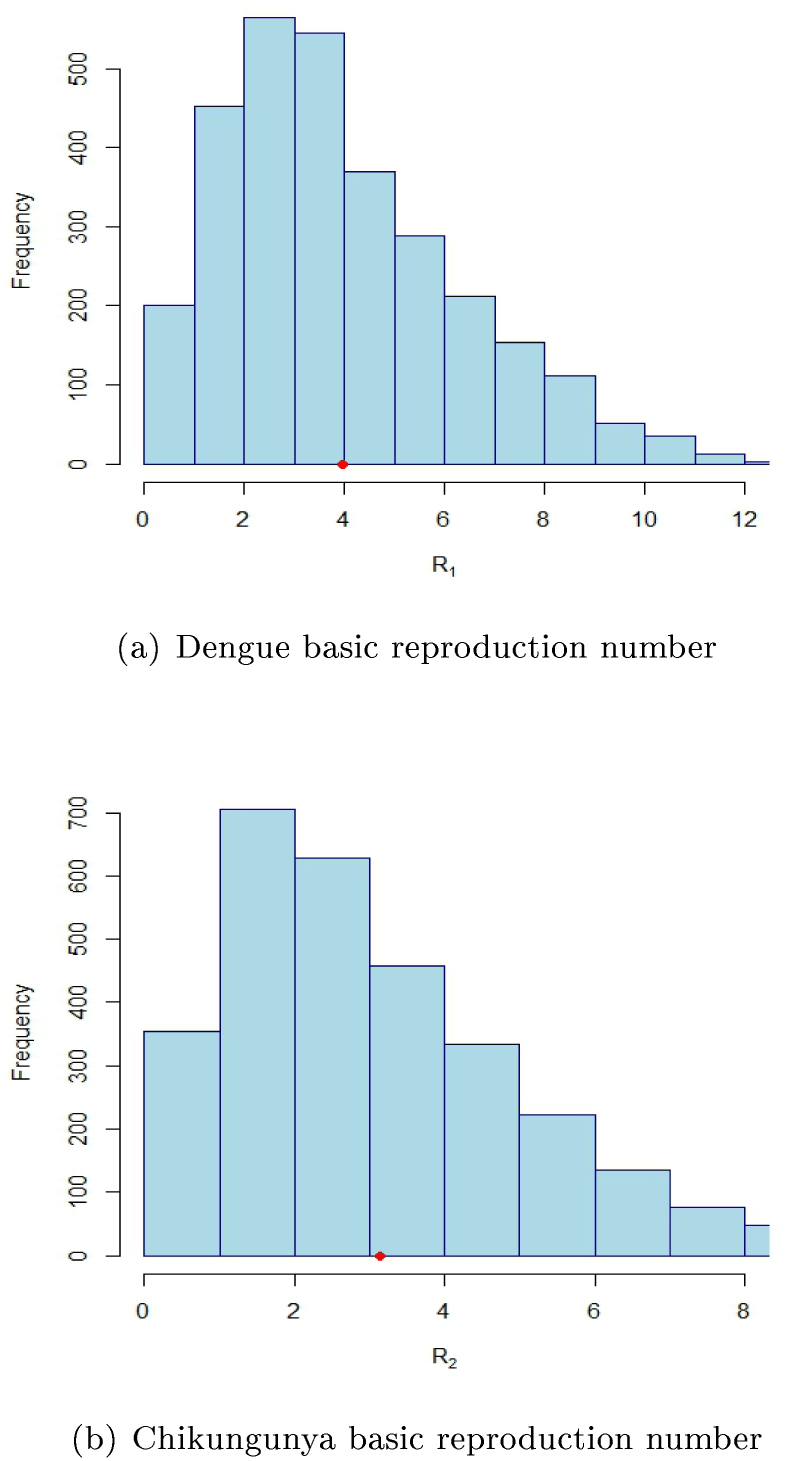
Histogram for Dengue and Chikungunya basic reproduction number. The basic reproduction number was obtained by 3000 Latin Hypercube Sampling simulations. The point in (a) and (b) indicates the mean of the sampling: *R*_1_ = 4.35 and *R*_2_ = 3.44, respectively.

With Latin Hypercube Sampling method we obtained parameter values, then we calculated partial rank correlation coefficients (PRCC), which determine the impact that each parameter has on the reproduction number, since this is a quantity of interes for us; moreover, this calculation takes into account the possible interplay between the remaining parameters. The parameters with PRCC values more closer to +1 or −1 are those who strongly influence on the outcome. The input and output variable keep a qualitative relationship given by the sign [21], see Fig. 11.

**Figure 11:**
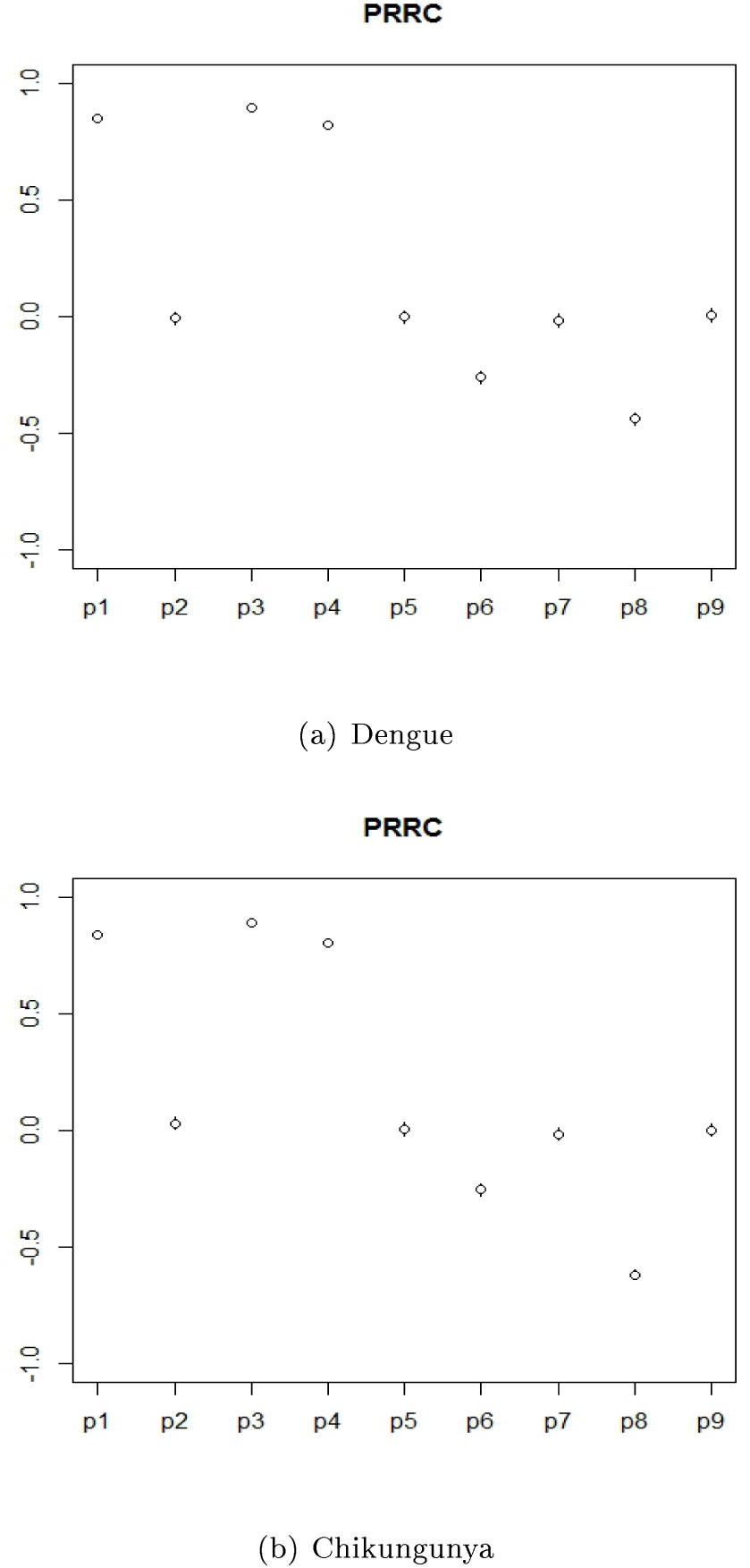
PRCCs for Dengue and Chikungunya basic reproduction numbers. We use Latin Hypercube Sampling method in each parameter ranges for the corresponding reproduction number, the relationship between the parameters and reproduction number is given by the sign, the parameters with magnitude closer to 1 have a greater effect on the reproduction number.

### 4.4 Effects of the components of the mosquito contact rates

In this section we show the effects on the diseases prevalence of the parameters that most affect to the basic reproduction numbers; these are components of the mosquito contact rate, for example remember that: *σ_υi_* is the number of bites that one mosquito would give a human, *β_hυi_* is the probability of virus transmission from an infectious mosquito to a susceptible human and *β_υhi_* is the probability of virus transmission from an infectious human to a susceptible mosquito (*i* = 1, 2).

According the Fig. 11 we note that when these parameters increase, they generate a higher basic reproduction number for both diseases, thereby we expect that a increase happen on the diseases prevalence.

For Fig. 12. *σ_hi_* = 19, *i* = 1,2, *σ_υ_*_2_ = 0.5, *β_υh_*_1_ = 0.35, *β_hυ_*_2_ = 0.55 and *β*_*υh*2_ = 0.54, we show that Dengue increases along with *β_hυ_*_1_ and it presents a major sensitivity to this parameter, on the other hand the Chikungunya prevalence decreases with respect to the same parameter.

**Figure 12:**
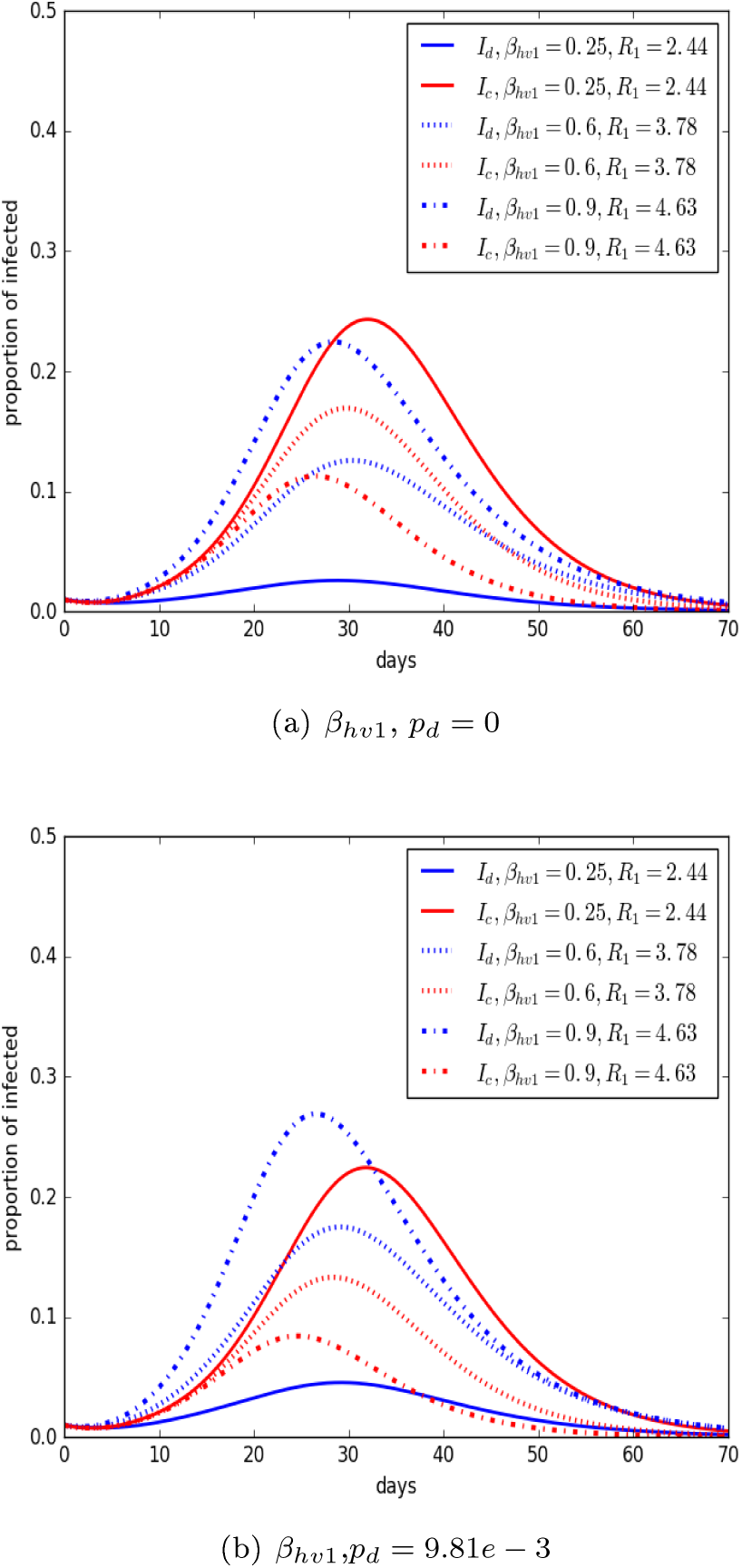
Plot of Dengue and Chikungunya prevalence corresponding to different scenarios to demonstrate the effects of changing *σ_υ_*_1_, (a) *p_d_* = 0 and (b) *p_d_* = 4.31*e* – 3 on the disease dynamics. The *y*-axis is the proportion of hots and the *x*-axis is time in days. We take *R*_2_ = 3.76 with parameters in Table 1.

Results for Chikungunya are shown in Fig. 13.

**Figure 13:**
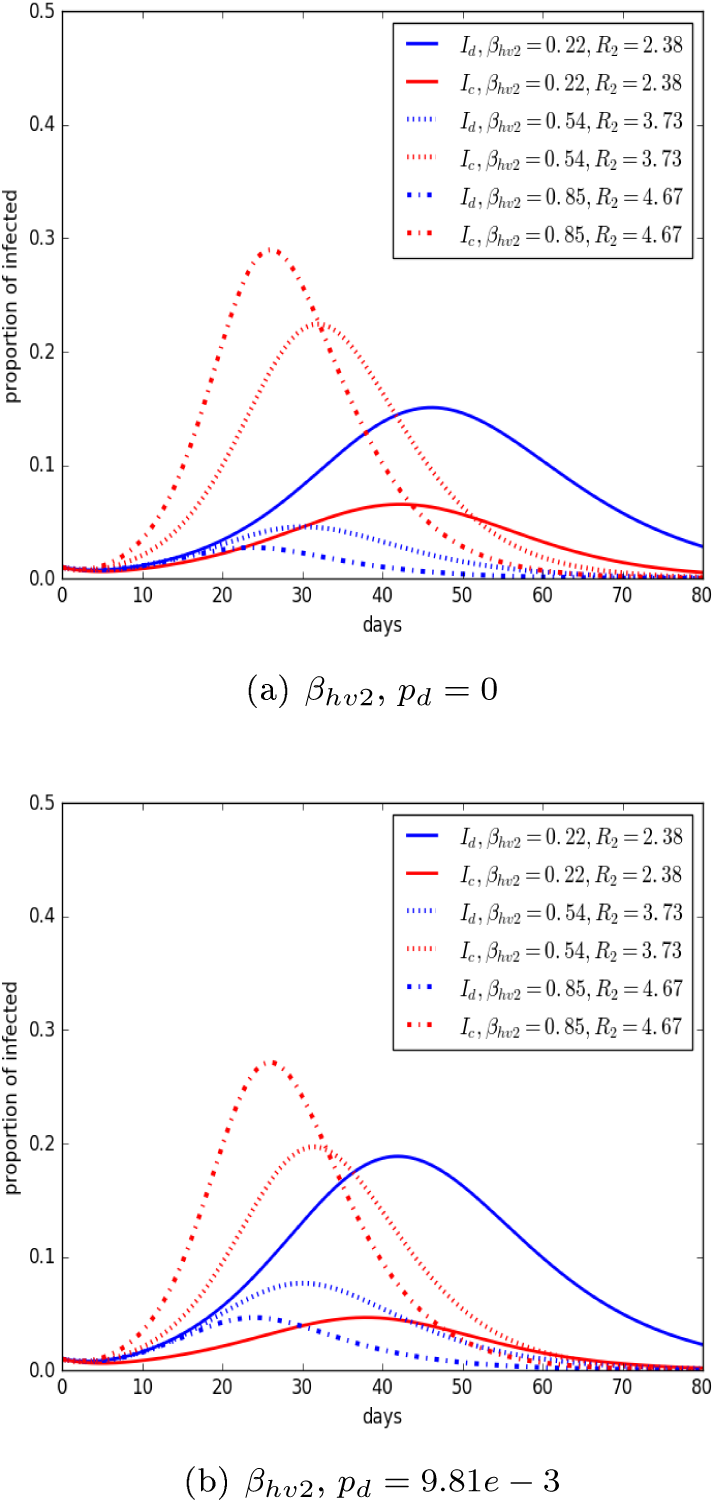
Plot of Dengue and Chikungunya prevalence corresponding to different scenarios to demonstrate the effects of changing *β_hυ_*_2_, (a) *p_d_* = 0 and (b) *p_d_* = 4.31*e* – 3 on the disease dynamics. The *y*-axis is the proportion of hots and the *x*- axis is time in days. We take *R*_1_ = 2.84 with parameters in Table 1

For Fig. 13. *σ_υi_* = 0.5, *σ_hi_* = 19, *i* = 1, 2, *β_hυ_*_1_ = 0.35, *β_υh_*_1_ = 0.34, and *β_υh_*_2_ = 0.54, we observe that the parameter increase along with *R*_2_ and the Chikungunya prevalence. We see a contrary trend on the Dengue prevalence.

In Fig. 14. We observe that when *γ*_1_ increases (i.e., number of days to recover from Dengue decrease), the Dengue basic reproduction number decreases. Also, we note that the Dengue prevalence decreases as *γ*_1_ increases, a contrary trend for the peak of Chikungunya prevalence.

**Figure 14:**
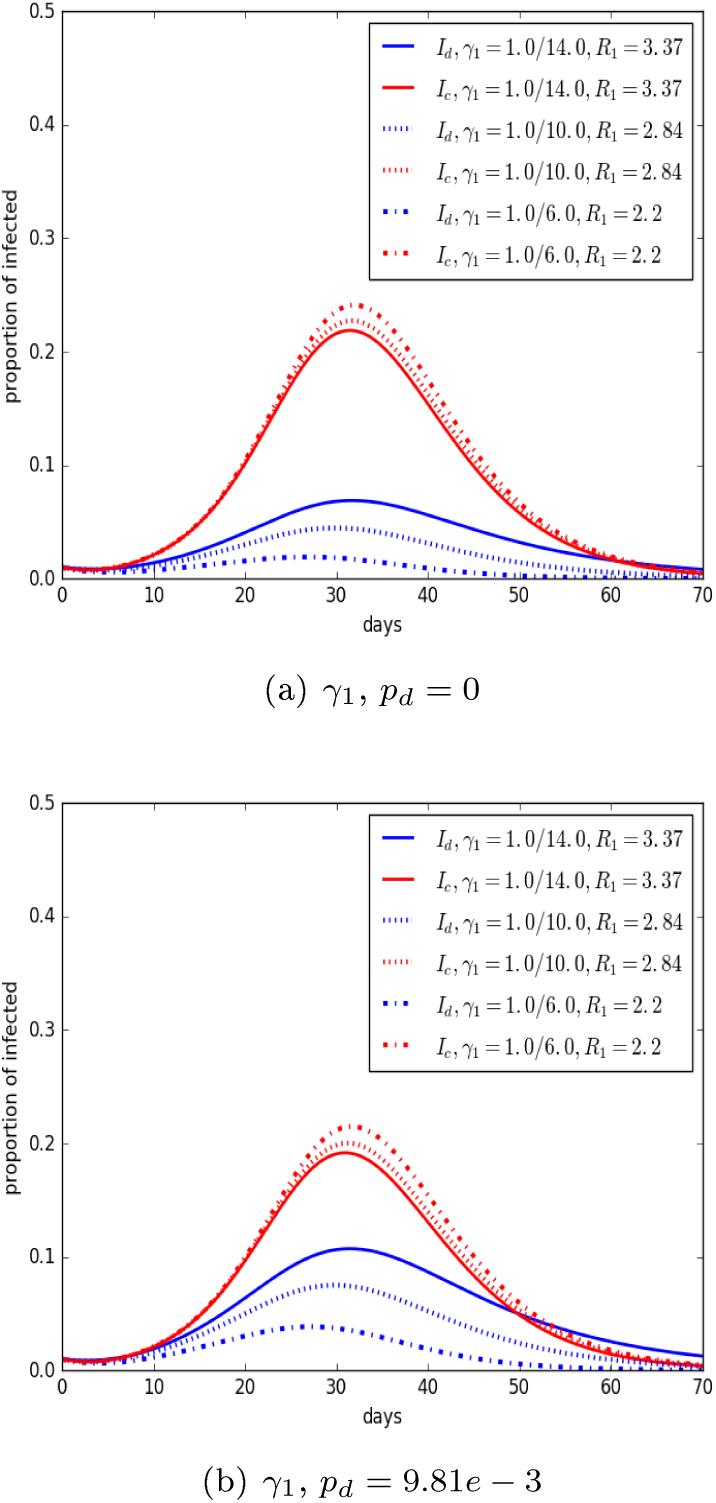
Plot of Dengue and Chikungunya prevalence corresponding to different scenarios to demonstrate the effects of changing *γ*_1_, (a) *p_d_* = 0 and (b) *p_d_* = 4.31*e* – 3 on the disease dynamics. The *y*-axis is the proportion of hots and the *x*-axis is is time in days. We take *R*_2_ = 3.76, with parameters in Table 1; moreover, *σ_υi_* = 0.5, *σ_hi_* = 19, *i* = 1, 2, *β_hυ_*_1_ = 0.35, *β_υh_*_1_ = 0.34, *β_hυ_*_2_ = 0.55 and *β*_*υh*2_ = 0.54.

In Fig. 15. We observe that when *γ*_2_ increases, the Chikungunya basic reproduction number decreases. Also, we note that the prevalence of Chikungunya decreases as *γ*_2_ increases and there are light changes in the infected of Dengue, but with a contrary trend to that one of Chikungunya.

**Figure 15:**
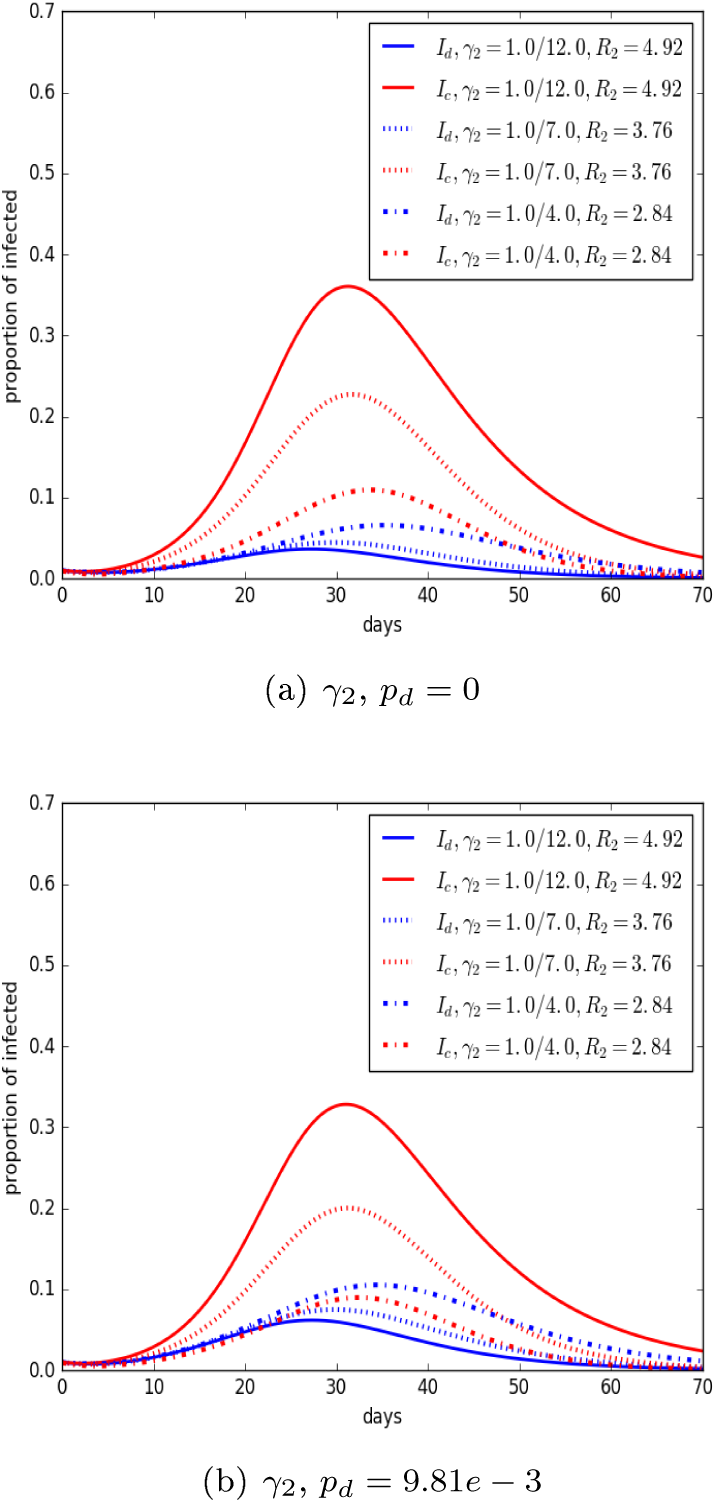
Plot of Dengue and Chikungunya prevalence corresponding to different scenarios to demonstrate the effects of changing *γ*_2_, (a) *p_d_* = 0 and (b) *p_d_* = 4.31*e* – 3 on the disease dynamics. The *y*-axis is the proportion of hots and the *x*-axis is is time in days. We take *R*1 = 2.84 with parameters in Table 1; moreover, *σ_υi_* = 0.5, *σ_hi_* = 19, *i* = 1, 2, *β_hυ_*_1_ = 0.35 *β_υh_*_1_ = 0.34, *β_hυ_*_2_ = 0.55 and *β_υh_*_2_ = 0.54.

In Fig. 16. We observe that when *δ* increases (i.e., the mosquito lifespan decreases), the Dengue and Chikungunya basic reproduction numbers decrease like their prevalences.

**Figure 16:**
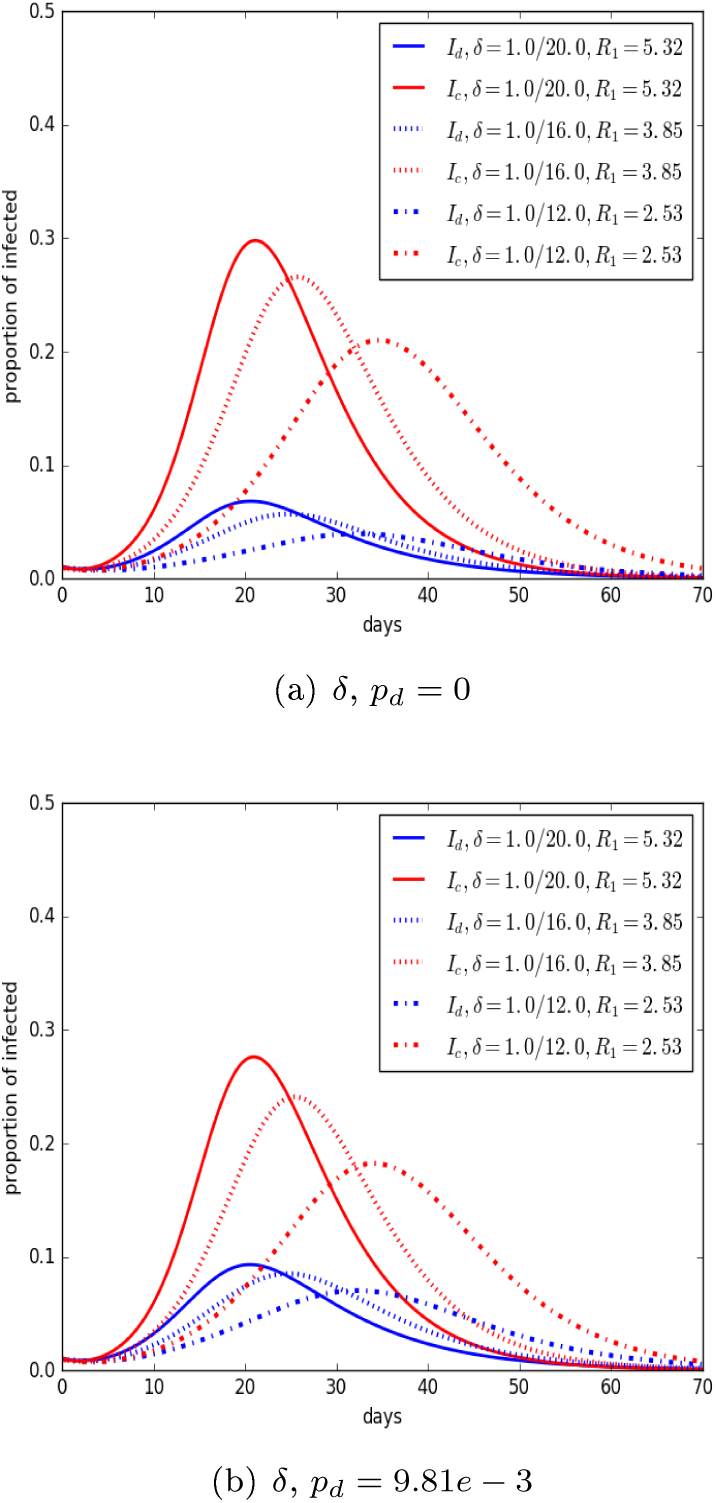
Plot of Dengue and Chikungunya prevalence corresponding to different scenarios to demonstrate the effects of changing *δ*, (a) *p_d_* = 0 and (*b*) *p_d_* = 4.31*e* – 3 on the disease dynamics. The *y*-axis is the proportion of hots and the *x*-axis is is time in days. We take *R*2 = 7.03, *R*_2_ = 5.09 and *R*_2_ = 3.34, respectively, with parameters in Table 1; moreover, *σ_υi_* = 0.5, *σ_hi_* = 19, *i* = 1, 2, *β_hυ_*_1_ = 0.35, *β_υh_*_1_ = 0.34, *β_hυ_*_2_ = 0.55 and *β_υh_*_2_ = 0.54.

**Figure 17:**
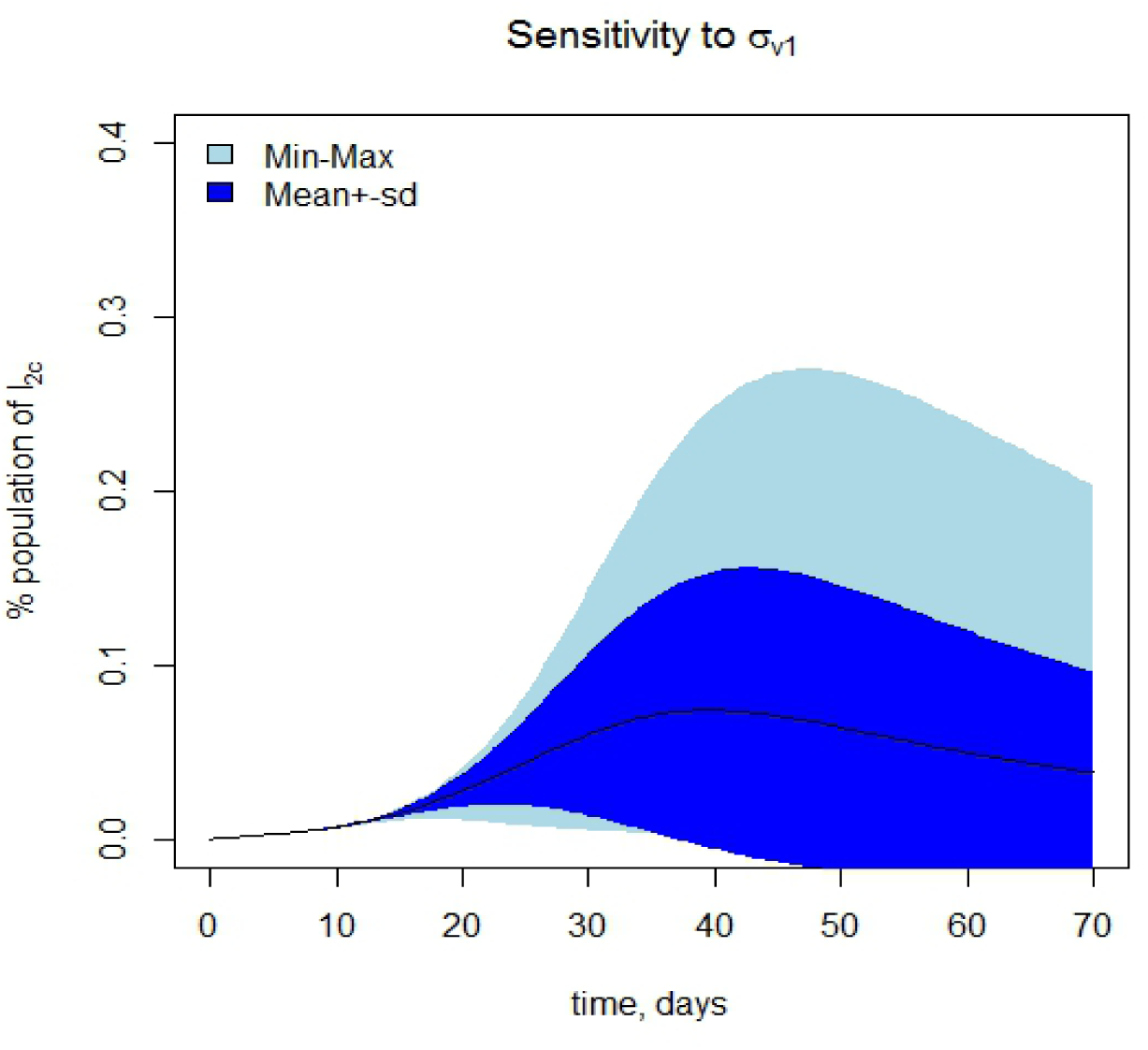
Sensitivity range for the parameter *σ_υ_*_1_. With min=0.33 and max=1.0, *σ_υ_*_2_ = 0.5 *σ_hi_* = 19, *i* = 1, 2 *β_hυ_*_1_ = 0.35 *β_υh_*_1_ = 0.34 and *β_hυ_*_2_ = 0.55, *β_υh_*_2_ = 0.54, so we will have *R*_1_ = 2.84 and *R*_2_ = 3.76, with parameters in Table 1. The model is run 1000 times, with Latin Hypercube Sampling algorithm. This is one of the most important parameter on *I*_2_*_c_*, the sub-acute state.

**Figure 18:**
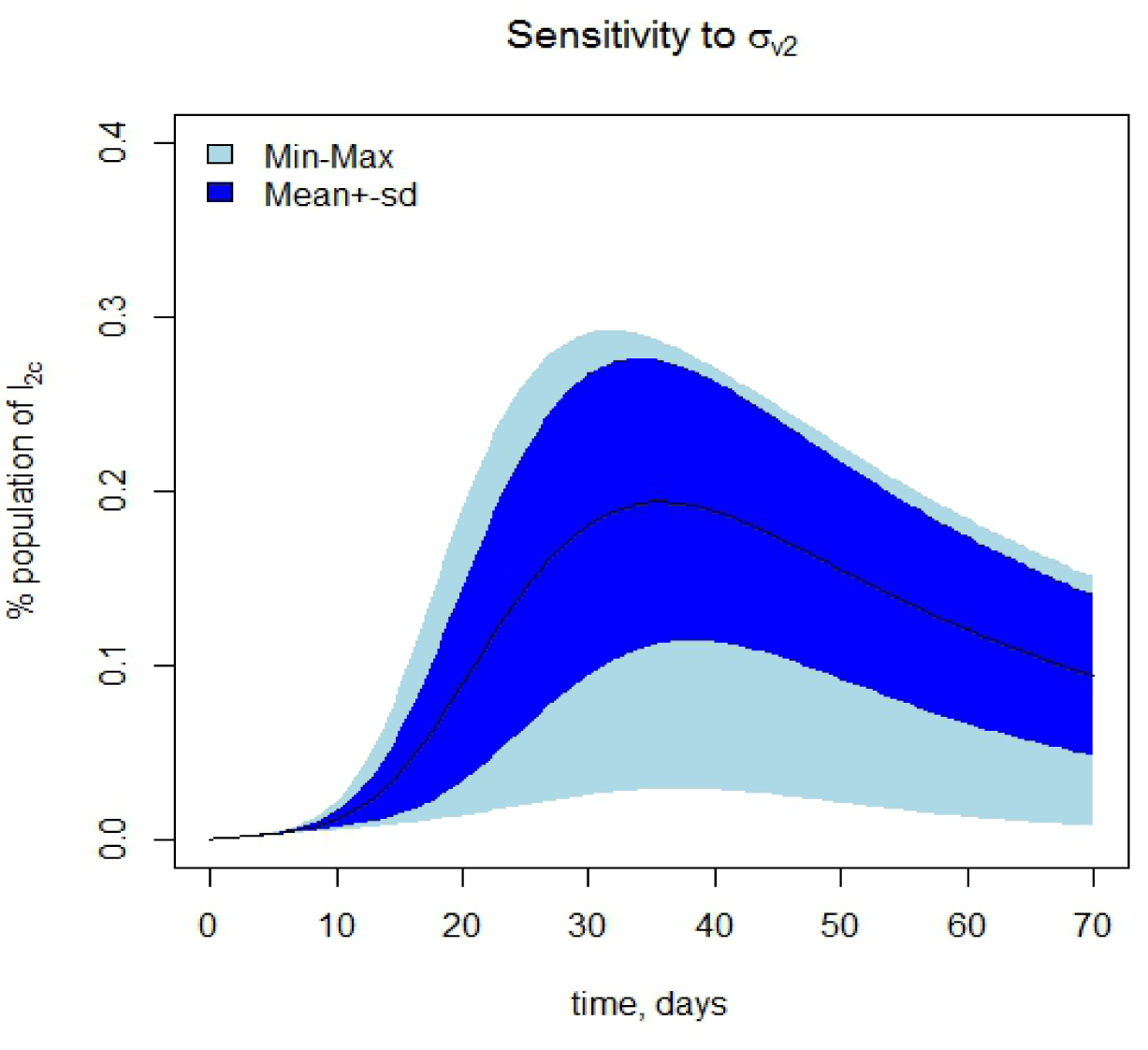
Sensitivity range for the parameter *σ_υ_*_2_. With min=0.33 and max=1.0, *σ_υ_*_1_ = 0.5 *σ_hi_* = 19, *i* = 1, 2, *β_hυ_*_1_ = 0.35 *β_υh_*_1_ = 0.34 and *β_hυ_*_2_ = 0.55, *β_υh_*_2_ = 0.54, so we will have *R*_1_ = 2.84 and *R*_2_ = 3.76, the parameters in Table 1. The model is run 1000 times, with Latin Hypercube Sampling algorithm.

**Figure 19:**
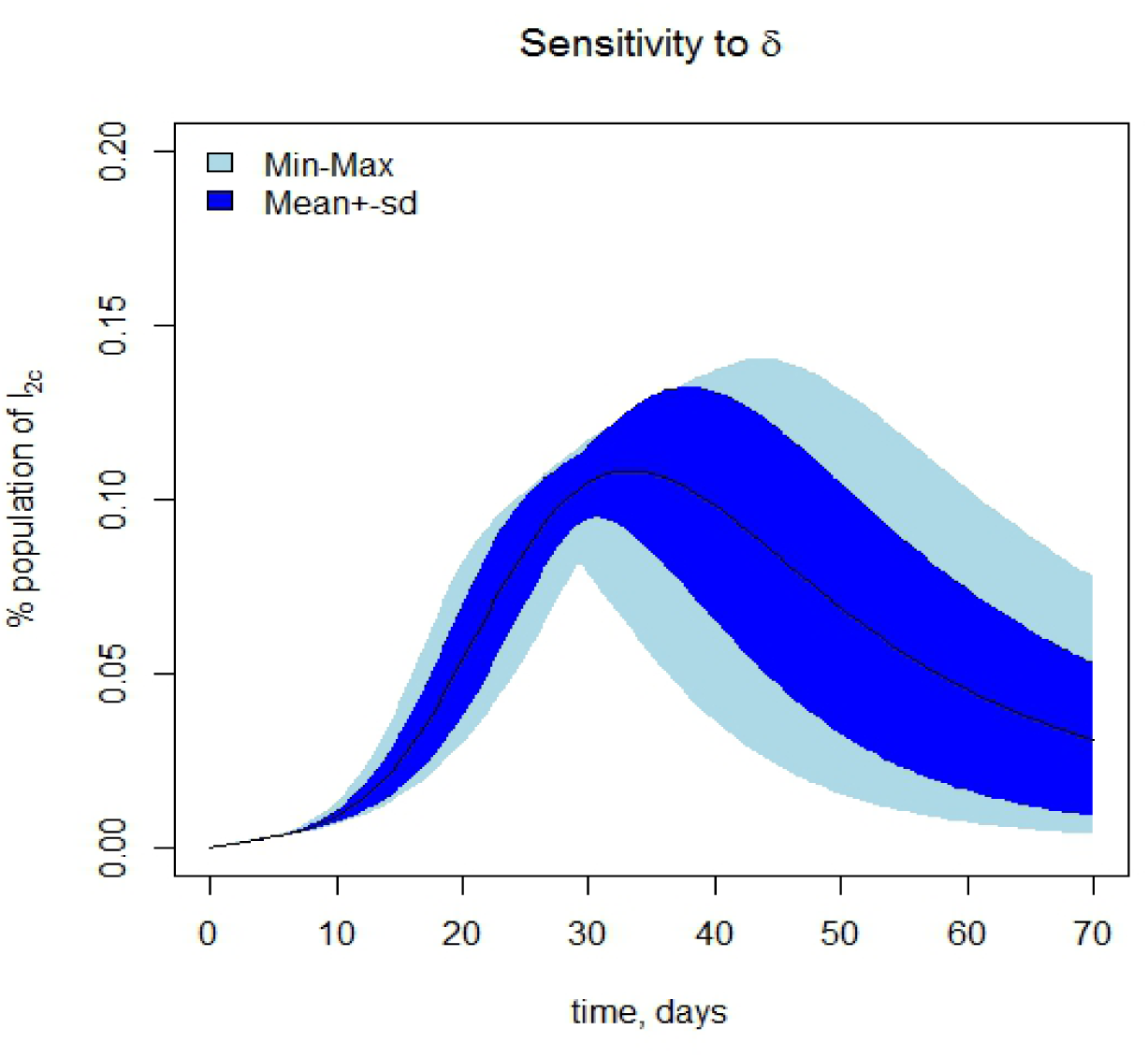
Sensitivity range for the parameter *δ*. With min=1.0/20.0 and max=1.0/12.0, *σ_υi_* = 0.5 *σ_hi_* = 19, *i* = 1, 2, *β_υh_*_1_ = 0.34 and *β_hυ_*_2_ = 0.55, *β_υh_*_2_ = 0.54, so we will have *R*_1_ = 2.84 and *R*_2_ = 3.76, with parameters in Table 1. The model is run 1000 times, with Latin Hypercube Sampling algorithm. This is other of the most important parameter on *I*_2_*_c_*, the sub-acute state.

### 4.5 Effects on sub-acute infected class (*I*_2*c*_)

We decided to explore the sensitivity to the model parameters that the *I*_2_*_c_* class presents for which we use R-package **FME**. We performed a local sensitivity analysis with the `sensFun` function, we estimated the effect that all the model parameters have on the variable *I*_2_*_c_*. This function generates some summaries, with which we can rank the parameters and determine those that most affect our variable of interest (see Table 11).

**Table 11:** Parameter ranking for the sensitivity of the *I*_2_*_c_* class. The more important parameter is that one with the higher absolute sensitivity value in L1 norm.

| Parameter | Sensitivity (L1) | Rank |
| --- | --- | --- |
| $\sigma_{v1}$ | 2.3e-01 | 1 |
| $\sigma_{h1}$ | 9.2e-03 | 17 |
| $\beta_{hv1}$ | 1.4e-01 | 4 |
| $\beta_{vh1}$ | 1.1e-01 | 8 |
| $\eta_1$ | 2.0e-02 | 16 |
| $\gamma_1$ | 5.6e-02 | 11 |
| $\mu$ | 1.4e-01 | 5 |
| $\sigma_{v2}$ | 1.9e-01 | 2 |
| $\sigma_{h2}$ | 7.6e-03 | 19 |
| $\beta_{hv2}$ | 9.5e-02 | 10 |
| $\beta_{vh2}$ | 1.0e-01 | 9 |
| $\eta_2$ | 3.1e-02 | 14 |
| $\gamma_2$ | 4.1e-02 | 13 |
| $\nu$ | 9.0e-03 | 18 |
| $\delta$ | 1.6e-01 | 3 |
| $\sigma_1$ | 4.5e-02 | 12 |
| $h$ | 1.4e-01 | 6 |
| $q$ | 1.4e-01 | 7 |
| $p_d$ | 2.7e-02 | 15 |

As the Table 11 shows that the most important parameters for the *I*_2_*_c_* class are: *σ_υ_*_1_, *σ_υ_*_2_ and *δ*, on the other hand, the parameter with less influence is *ν*. So with this in mind, we performed a global sensitivity analysis, where the parameters were varied over the ranges in the Table 7 and 8, and the effect on *I*_2_*_c_* was assessed. From the R-package **FME** we use the function `sensRange`. We define one or more sensitivity parameters and we assign a distribution; then we have to stipulate the minimum and maximum values that each parameter can take. In our case we decide to estimate the sensitivity that present the variable *I*_2_*_c_* when we choose only one sensitivity parameter, for example, what happen if we vary the values of *δ* according to Latin Hypercube Sampling algorithm. To do this, the model is run 1000 times.

## 5 Discussion

In this paper, we have constructed and explored the coupled dynamics of Dengue and Chikungunya when they invade the same host population in the presence of only one vector species *Ae. aegypti*, that transmits both viruses.

We first calculate the basic reproduction number for each disease, using the Latin Hypercube Sampling to find the mean of each basic reproduction number. Then we compare the results of the model with Dengue and Chikungunya data from Mexico 2015, 2013 and we obtain a good fit between both, this means that our model has the capacity to predict the fate of the diseases. We also performed numerical simulations to examine the population dynamics under various scenarios to determine the effect of the basic reproduction numbers on prevalences; moreover, we analyze the effect of infection parameters on the susceptible population, resulting that *σ_υi_* (number of bites that one mosquito would give a human per unit time) increases while susceptible population decreases. Our model predicts the coexistence of both viruses with approximately the same outbreak period. We performed a sensitivity analysis to determine which parameters influence to the reproduction numbers and *I*_2_*_c_* class (sub-acute phase of Chikungunya); we chose this class because the infected individuals in this phase only can pass to a chronic phase without recovery along with a greater susceptibility to Dengue infection than the other categories.

The study of Chikungunya has risen since it is a reemerging disease. Cauchemez et. al. [25] describe the spread of Chikungunya virus (CHIKV) in the first three areas (Saint Martin, Martinique and Guadeloupe) of the Caribbean. They highlight the effectiveness of local transmission of CHIKV, and report the number of new cases that an infected human can generate: 2-4, which roughly coincides with our results since our estimates for the Chikungunya basic reproduction number has a mean of 3.44 (see Fig. 10(b)).

*R*_0_ values estimated by Cauchemez et. al. [25] and by us, indicate that there may be a spatial homogeneity among the *Ae. aegypti* mosquito population and the susceptible population to CHIKV infection, that promotes outbreaks and allows a permanent high prevalence. This spatial homogeneity may be due to the degree of disability inflected on the infected subjects, that remain exposed to mosquito bites thus increasing the vectorial capacity of *Ae. aegypti*. In the case of Dengue, the clinical symptoms are mild, allowing people to move even in the infective phase to places where the densities of the vectors can be varied and thus increases the spatial heterogeneity in the transmission. Records of attack rates in the Americas indicate that all age groups are susceptible to infection, reaching an overall attack rate of 80%, which confirms that *R*_0_ may be similar in places where CHIKV is being introduced. Padmanabha et al. [26] found *Ae. aegypti* rests, feeds and lays eggs near homes, in close physical proximity to humans. Consequently, the females *Ae. aegypti* feed on humans repeatedly.

For Dengue, Nishiura [20] did a compilation of Dengue results from mathematical and statistical proposals, among the main results is the calculation of *R*_0_, our estimate agrees with these results for the Americas with a mean of 4.35.

On the other hand, Pandey et. al. [24] performed uncertainty and sensitivity analysis of *R*_0_ for their model using partial rank correlation coefficients (PRCC) and concluded (Fig. 5. in [24]) that the more influential parameters on *R_0_* are the transmission rates (with positive influence) and the average lifespan of a mosquito along with the human recovery rate with negative influence (see Fig. 11. for our results).

With respect to the sensitivity of *R*_0_ we show that the most sensitive parameters are: the mosquito biting rate and the transmission rates for both diseases (positive influence) and the average lifespan of a mosquito along with the human recovery rate, with a negative influence for each disease. Our results (Fig. 11) are consistent with [1]. Acevedo et al. [27] have shown that low mobility can eventually result in homogeneity in the values of *R*_0_, spatial heterogeneity in the population of *Aedes* mosquitoes and the human population. The presence of Chikungunya virus in Mexico was recorded with around of 155 cases and 11500 cases for 2014, 2015 respectively, throughout the country; which indicate both a broad attack rate and spread of the disease in susceptible people.

Besides the effect that had the parameters on *R_i_, i* = 1, 2, we also analyzed the effect on other variables. We observed that the transmission parameters *σ_υi_* (number of bites that one mosquito would give a human per unit time), *σ_hi_* (maximum number of bites that a human can suffer per unit of time), *β_hυi_* (probability of virus transmission from an infectious mosquito to a susceptible human), *β_υhi_* (probability of virus transmission from an infectious human to a susceptible mosquito), *i* = 1, 2, have a positive influence on the basic reproduction numbers, as well as on the infected populations of both diseases. Nevertheless this positive effect is observed more clearly with *β_hυ_*_1_ for Dengue and with *β_hυ_*_2_ for Chikungunya virus.

Our numerical results suggest that the application of typical strategies of vector control could help decreasing the corresponding disease prevalence since *σ_υi_* and *σ_hi_*, *i* = 1,2 will decrease by reducing vector population, host exposure, respectively [4]; but, at the same time this means that the other virus will increase, so it is necessary to set strategies to reduce both infected populations.

On the other hand, *δ* (mosquito lifespan) has a reverse effect on the basic reproduction numbers, this means that when *δ* increases each *R_i_*,*i* = 1,2, decreases, together with the infected populations of Dengue and Chikungunya. This is consistent with health strategies that focus on the eradication of vector population [4].

The parameter *γ_i_* (recovery rate), *i* = 1,2, has a reverse effect on the basic reproduction number *R_i_*, also the infected population with the corresponding subscript decreases, but in contrast, we note that the other infected population increases. This result indicates that strategies to reduce transmission days of a specific disease, at the same time they can help to increase infected cases of the other co-circulate virus.

Finally, we perform a sensitivity analysis for *I*_2_*_c_* (sub-acute phase of Chikungunya) and we found the most important parameters that influence it: *σ_υ_*_1_, *σ_υ_*_2_, *δ*; on the other hand, the parameter with less influence is *ν*, namely, *I*_2_*_c_* is primarily affected by changes in mosquitoes bite rates and the mosquito lifespan.

The work presented here is relevant for the introduction of Dengue and Chikungunya strains into novel areas, which is timely in the light of the spread of Chikungunya throughout the Americas, specifically into Mexico. We have a model that involving two co-circulating viruses, unlike the aforementioned models that only study the viruses for separate, we calculated a explicit formula for both basic reproduction numbers, we have analyzed different scenarios for the most important parameters in the spread of these diseases, we hope that this will help to the appropriate institutions for their control and eradication, in this regard, we recommend to implement a joint treatment (both recovery rates must be considered) in order to the infected population of any of the two diseases does not increase more than the other; moreover: the mosquito control (via insecticides), decrease the human exposition to mosquito bites via mosquito nets, do not staying in open spaces for a long time, etc., will help to reduce both diseases prevalence too if appropriate strategies are set. In later works we could do a detailed differentiation and analysis between four Dengue serotypes and seasonality.

## Acknowledgements

Jorge X. Velasco-HernÃąndez acknowledges support from grant PAPIIT UNAM IN110917. Additional thanks to Dr. Brenda Tapia Santos for her thoughtful comments and observations. Nohemí Báez-Hernández acknowledges the support from CONACyT through the scholarship No. 245051 and FORDECyT.

